# Chronic kidney disease promotes anxiety susceptibility through an angiotensin II–central amygdala axis

**DOI:** 10.64898/2026.08.16.744184

**Authors:** Yuting Liu, Yujian He, Xinzhou Zhang, Zhen Wang, Lu Zhang, Nan Hu, Hualin Ma, Fan Yang

**Author notes:** Corresponding authors: Fan Yang, PhD. Shenzhen Institute of Advanced Technology,; Hualin Ma, PhD. Shenzhen People’s Hospital,; Nan Hu, PhD. Shenzhen People’s Hospital,; Lu Zhang, PhD. Shenzhen Institute of Advanced Technology. These authors contributed equally to this work (Yuting Liu, Yujian He).

## Abstract

**Background:** Neuropsychiatric comorbidities are highly prevalent in chronic kidney disease (CKD), yet the underlying neural mechanisms remain poorly defined.

**Methods:** We established multiple mouse models of CKD and identified an adenine-induced model as the most suitable platform to study neurobehavioral alterations. Anxiety susceptibility was operationalized as the emergence of anxiety-like behavior after subthreshold unpredictable stress (SUS) and was assessed using the SUS paradigm combined with behavioral assays. Region-focused c-Fos mapping, fiber photometry, and chemogenetic manipulation were used to interrogate neural circuit activity. Pharmacological and genetic approaches were applied to investigate the role of angiotensin II (Ang II) signaling. Finally, hypothalamic paraventricular nucleus (PVN) activation was used to explore brain-to-kidney feedback by using in vivo multiphoton microscopy imaging techniques.

**Results:** CKD mice showed no consistent baseline anxiety-like phenotype across standard assays but developed robust anxiety-like behavior after subthreshold unpredictable stress. Region-focused c-Fos profiling and fiber photometry identified the central amygdala (CeA) as a stress-sensitized limbic node in CKD. Chemogenetic inhibition of CeA GABAergic neurons attenuated anxiety-like behavior, supporting a functional role for CeA activity. Mechanistically, CKD elevated circulating Ang II and enhanced CeA accumulation of peripherally administered FAM-Ang II-associated signal. CeA-specific *Agtr1a* knockdown attenuated anxiety-like behavior and exaggerated stress-evoked CeA calcium responses. Exploratory experiments further showed that sustained PVN glutamatergic activation aggravated early renal injury markers in a mild renal injury model. These findings support a kidney-to-brain model in which CKD primes CeA stress circuits, while local Ang II–AT1R signaling contributes to the behavioral expression of stress-induced anxiety-like behavior, with a potential brain-to-kidney feedback component.

**Conclusions:** CKD promotes stress-induced anxiety susceptibility through a CeA-centered mechanism involving local Ang II–AT1R signaling. These findings identify CeA Ang II–AT1R signaling as a potential contributor to CKD-associated stress-related affective vulnerability.

**Abstract graph:** 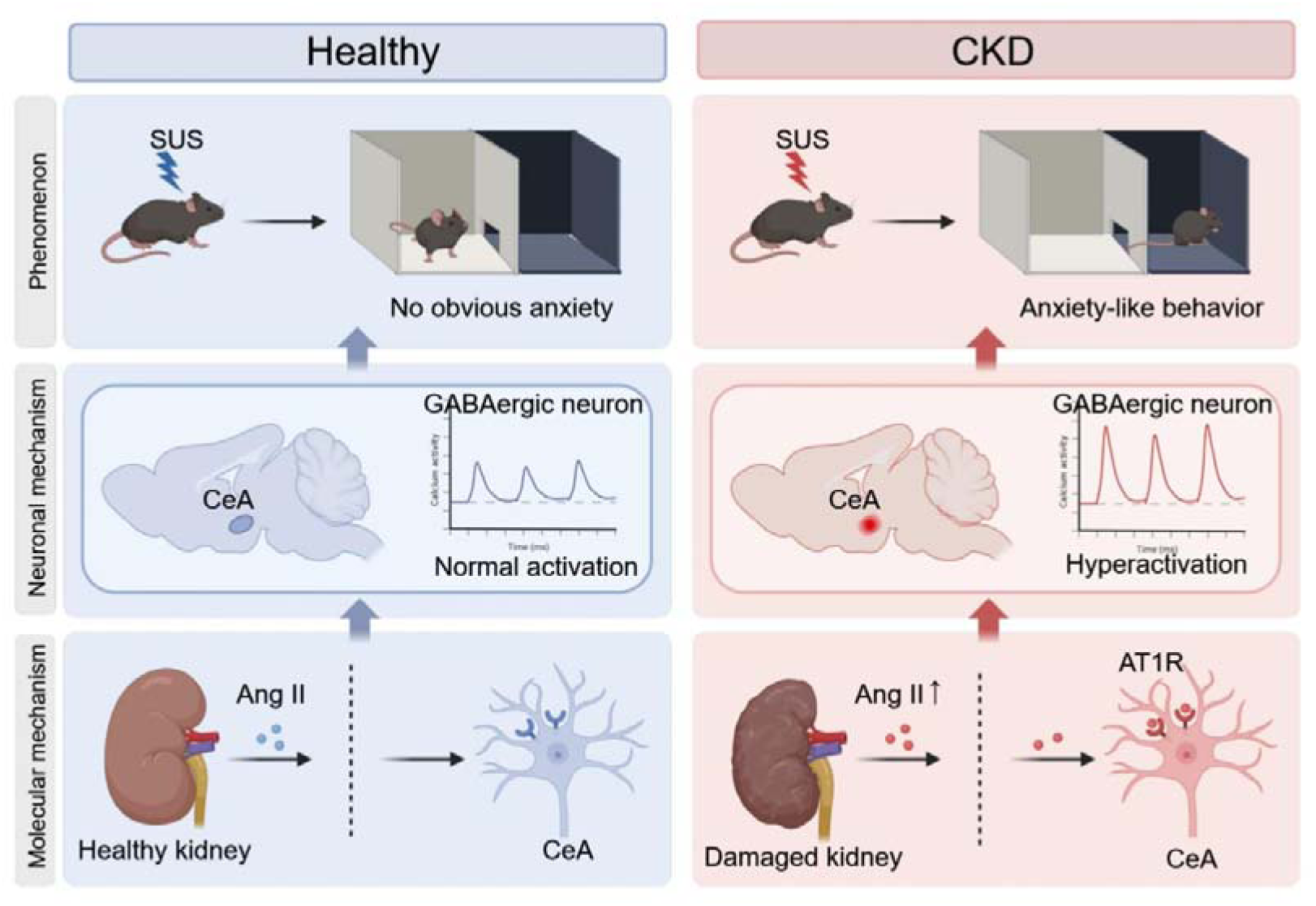

## Introduction

Chronic kidney disease (CKD) affects approximately 10% of the global adult population and is a leading cause of morbidity and premature mortality worldwide^[1]^. Beyond its renal and cardiovascular consequences, CKD is associated with a high prevalence of neuropsychiatric comorbidities, particularly anxiety and depression, which substantially impair quality of life and worsen long-term prognosis^[2,3]^. These affective disturbances are especially prominent in patients with advanced or end-stage disease, yet their underlying neurobiological mechanisms remain poorly understood, and effective targeted interventions are lacking^[4]^. CKD is increasingly recognized as a systemic condition that disrupts bidirectional communication between the kidney and the brain.

Accumulating evidence indicates that uremic toxins, chronic inflammation, oxidative stress, and blood–brain barrier (BBB) disruption collectively impair central nervous system homeostasis and may predispose patients to affective dysregulation^[5,6]^. Clinical neuroimaging studies have provided direct support for central nervous system involvement in CKD: patients with end-stage renal disease exhibit reduced gray matter volume in limbic regions and weakened functional connectivity between the amygdala and prefrontal or cingulate cortices—changes that correlate with emotional symptom burden^[7,8]^. These findings implicate the amygdala as a key node in CKD-related affective pathology, but the resolution of conventional fMRI precludes identification of which amygdalar subregion drives this phenotype.

Among the systemic pathways that may link renal dysfunction to central circuit alterations, the renin–angiotensin system (RAS) is a compelling candidate. Intrarenal RAS dysregulation can occur early during kidney disease and contribute to renal injury before overt deterioration of filtration function in some CKD settings^[9]^.The brain also contains a functionally active angiotensin signaling system that regulates neurotransmission, cerebrovascular function, stress responsiveness, and emotional behavior^[10,11]^. Because CKD is associated with impaired BBB integrity, sustained elevation of circulating Ang II may increase the engagement of AT1R-expressing brain regions and perturb local angiotensin signaling. However, whether peripheral Ang II engages a specific amygdalar subregion to promote stress-induced anxiety-like behavior in CKD remains unknown. Moreover, because psychiatric comorbidities in chronic disease may manifest as exaggerated responses to mild stressors rather than overt baseline abnormalities, standard behavioral assessment under resting conditions may fail to capture CKD-associated affective vulnerability—a consideration that has been largely overlooked in preclinical CKD research^[12,13]^. Accordingly, in this study, anxiety susceptibility was operationalized as the emergence of anxiety-like behavior after subthreshold unpredictable stress (SUS), rather than as a stable baseline anxiety-like phenotype.

In this study, c-Fos mapping identified the CeA and PVN as candidate stress-related regions in the adenine-induced CKD model. The CeA was prioritized for mechanistic investigation based on its significant CKD-associated activation and established role in anxiety-related behavior. Using fiber photometry, chemogenetic manipulation, and local *Agtr1a* knockdown, we found that CeA Agtr1a signaling contributes to stress-associated CeA hyperresponsiveness and anxiety-like behavior. Exploratory experiments further showed that sustained experimental activation of PVN glutamatergic neurons increased selected renal injury markers, providing preliminary evidence consistent with a potential brain-to-kidney feedback component. Together, these findings define a CeA–Ang II axis as a previously unrecognized mechanism linking renal dysfunction to central affective dysregulation, and suggest that stress-responsive central circuits may in turn amplify kidney injury in the context of CKD.

## Methods

### Animals

Male C57BL/6N mice, 8–10 weeks of age and weighing 25–30 g, were obtained from Guangdong Vital River Laboratory Animal Technology (Foshan, China) and maintained in the specific pathogen-free animal facility at XXXX. Gad2-IRES-Cre mice (JAX stock no. 010802) were originally purchased from The Jackson Laboratory and maintained on a C57BL/6J background in the animal facility at XXXX. Experimental male animals, aged 8–10 weeks, were generated by in-house breeding. Animals were housed in a temperature-controlled room at 20–25 °C under a 12-h light/dark cycle, with free access to food and water unless otherwise indicated. Mice were allowed to acclimatize for at least 1 week before experimental procedures.

All procedures were carried out in accordance with the protocols approved by the Ethics Committee of the Animal Care and Use Committee of XXXX (AUP-240417-LYT-256-01) and XXXX (SIAT-IACUC-190219-NS-YF-A0582). Animals were randomly assigned to experimental groups. Behavioral testing, histological quantification and imaging analyses were performed by investigators blinded to group allocation whenever possible.

### Chronic kidney disease models

Three chronic kidney injury models were generated in male C57BL/6N mice to identify a platform suitable for studying CKD-associated behavioral alterations. Renal ischemia–reperfusion injury (IRI) was performed under isoflurane anesthesia (3–4% induction, 1–2% maintenance in O_2_) . Bilateral renal pedicles were exposed through dorsal flank incisions and occluded using atraumatic microvascular clamps for 35 min^[14]^. Successful ischemia was confirmed by darkening of the kidney, and reperfusion was verified by restoration of renal color after clip removal.

The 5/6 nephrectomy model was established using a two-stage ligation-based procedure^[15]^. In the first surgery, the upper and lower poles of the left kidney were ligated with 4-0 suture. One week later, the right kidney was removed after renal pedicle ligation. Mice were analyzed 6 weeks after the second surgery.

For the two surgical CKD models, a common sham-operated cohort was used during the model-screening phase to reduce animal use in line with the 3R principle. Sham mice underwent renal exposure without vascular occlusion, ligation or parenchymal removal and served as the surgical control for both IRI and 5/6 nephrectomy comparisons.

For adenine-induced CKD, mice were fed a custom diet containing 0.25% adenine (Pythonbio, Cat. No. SY19004, Guangzhou Peiyu Biological Products Co., Ltd., Guangzhou, China) for 3 weeks^[16]^. Control mice received standard chow. After model comparison, the adenine-induced model was used for subsequent mechanistic experiments and is referred to as the CKD model. For mild renal injury experiments, mice were fed the same 0.25% adenine diet for 1 week.

### Enzyme-linked immunosorbent assay (ELISA)

Serum, cerebrospinal fluid (CSF) and urine samples were collected for ELISA-based measurements. Peripheral blood was collected from mice deeply anesthetized with sodium pentobarbital (100 mg/kg, *i.p.*) . Serum was isolated by centrifugation and stored at −80 °C until analysis. CSF was collected from the cisterna magna under isoflurane anesthesia (3–4% induction, 1–2% maintenance in O_2_) , and samples visibly contaminated with blood were discarded. When CSF samples were pooled because of limited material, each pool, rather than each individual mouse, was treated as one biological replicate. Urine was collected by gentle suprapubic stimulation and kept on ice before storage.

Serum and CSF Ang II concentrations were measured using a Mouse Angiotensin II ELISA Kit (RayBiotech, Cat. No. EIA-ANGII, Guangzhou, China). Urinary microalbumin (MAU), serum kidney injury molecule-1 (KIM-1) and neutrophil gelatinase-associated lipocalin (NGAL), were measured using commercial ELISA kits (Shanghai Enzyme-linked Biotechnology, Cat. No. ml037573A, ml002075A, ml002141A, Shanghai, China). Urinary albumin was normalized to urinary creatinine and expressed as the albumin-to-creatinine ratio (UACR). The UACR was calculated using the following formula: UACR (mg/g) = urinary albumin (mg/L) / urinary creatinine (g/L). All assays were performed according to the manufacturers’ instructions. Absorbance was measured using a microplate reader (Infinite 200 PRO, Tecan, Switzerland).

### Renal biochemical assessment

Serum creatinine (Scr) and blood urea nitrogen (BUN) levels were determined using a Creatinine Assay Kit (Nanjing Jiancheng Bioengineering Institute, Cat. No. C011-2-1, Nanjing, China), and a Urea Assay Kit (Nanjing Jiancheng Bioengineering Institute, Cat. No. C013-2-1, Nanjing, China), respectively, according to the manufacturer’s instructions.

### Renal histology

Mice were deeply anesthetized with sodium pentobarbital (100 mg/kg, *i.p.*) and transcardially perfused with phosphate-buffered saline (PBS) followed by 4% paraformaldehyde (PFA). Kidneys were collected, post-fixed in 4% PFA for at least 24 h, dehydrated, embedded in paraffin and sectioned at 4 μm using a room-temperature microtome (RM2235, Leica, Germany).

Sections were stained with hematoxylin and eosin (H&E), Masson’s trichrome or periodic acid–Schiff (PAS) staining following standard protocols ^[17]^. Images were acquired using a slide scanner (VS200-BU, Olympus, Japan). Renal histopathological analysis was performed according to established protocols ^[18]^. Tubular injury was scored on H&E sections using a 0-4 semi-quantitative scale based on tubular dilation, atrophy, epithelial degeneration, casts, interstitial edema, fibrosis and inflammatory infiltration. Interstitial fibrosis was quantified on Masson-stained sections as the percentage of collagen-positive area using ImageJ. Glomerular injury was evaluated on PAS-stained sections from 20 randomly selected glomeruli per mouse using a 0-4 scale based on mesangial expansion, basement membrane thickening and sclerosis.

### Subthreshold unpredictable stress (SUS) model

A modified SUS paradigm was used to assess anxiety-like behavior after mild stress exposure, as previously described with minor modifications^[19]^. This paradigm was designed to be milder than conventional chronic unpredictable mild stress procedures and did not induce overt behavioral abnormalities in healthy mice. SUS exposure was initiated after the completion of adenine diet-induced CKD modeling, during which adenine-containing diet was discontinued to minimize the potential influence of residual adenine exposure on behavioral outcomes. Briefly, mice were exposed to one mild stressor per day for 10 consecutive days, and the same stressor was not applied on two consecutive days. The stressors included wet bedding, crowding, brief tail suspension,overnight illumination and daytime darkness. Behavioral assays were performed after the completion of stress exposure.

### Behavioral tests

Adenine-containing diet was discontinued at least 1 week before behavioral testing and replaced with the same standard chow used for control mice. Mice were transferred to the testing room 1–2 h before each experiment for habituation. Deschloroclozapine (DCZ, 6 μg/kg, MedChemExpress, HY-42110, USA) was injected 30 min before the chemogenetic manipulation experiment to inhibit neurons.

Tests were performed in the following order: anxiety-like behavioral assays, including the open-field test (OFT), elevated plus maze (EPM) and light–dark box (LDB) test; tail-suspension test (TST); and fear conditioning. Apparatuses were cleaned with 20% ethanol between animals to remove odor cues.

### OFT

The OFT was used to assess anxiety-like behavior. Mice were individually placed in a 50 × 50 cm square arena and allowed to explore freely for 10 min after 1 min of acclimation. The central zone was defined as the inner 25 × 25 cm area. Center entries, distance traveled in the center zone and total distance were quantified using TopScan Lite software (CleverSys, USA) .

### EPM

The EPM consisted of two open arms and two closed arms arranged in a cross configuration (arm size 25 × 5 cm, height 65 cm). Each mouse was placed on the central platform and recorded for 5 min after 1 min of acclimation using an overhead-mounted camera. Video were analyzed using EthoVision XT (Noldus, Netherlands) to quantify time spent in the open arms, as indices of anxiety-like behavior.

### LDB

The LDB contained two equal compartments connected by a small doorway. Mice were placed in the center of the illuminated compartment and allowed to explore freely for 5 min after 1 min of acclimation.

Light-chamber entries and time spent in the light compartment were quantified using EthoVision XT (Noldus, Netherlands), as indices of anxiety-like behavior.

### TST

Depression-like behavior was assessed using an automated tail-suspension system (BIO-TST, BioSeb, France). Mice were suspended by the tail for 6 min, and immobility time was recorded as the primary outcome.

### Fear conditioning

Fear conditioning was used to assess acute fear responses and fear-memory recall. The test was performed using a FreezeScan system (CleverSys, USA). Mice were habituated to the conditioning chamber for 5 min one day before training. On the training day, animals were placed in context A for 5 min and then exposed to five tone–shock pairings. Each conditioned stimulus consisted of a 30-s tone at 6,000 Hz and 80 dB, co-terminating with a 1-s foot shock at 0.5 mA. Pairings were separated by 120-s intervals. On the following two days, mice were tested in a modified context B and exposed to tone-only presentations to evaluate cued fear-memory recall.

Freezing was quantified as the percentage of time spent immobile.

### Metabolic monitoring

Circadian activity and respiratory metabolism were measured using an integrated metabolic monitoring system (Oxymax/CLAMS-HC, Columbus Instruments, Columbus, USA). Mice were individually housed in metabolic cages and acclimatized before recording. Oxygen consumption, carbon dioxide production, and wheel-running activity were continuously monitored over a 24-h period under the same light/dark cycle as the home cage. Food intake and water intake were recorded during the same metabolic cage monitoring period and expressed as grams per 24 h. Respiratory exchange ratio (RER) was calculated as VCO /VO.

### Histological analysis of brain tissue

Mice were deeply anesthetized with sodium pentobarbital (100 mg/kg, *i.p.*) and transcardially perfused with PBS followed by 4% PFA. Brains were removed, post-fixed in 4% PFA at 4 °C overnight, then transferred to 30% w/v sucrose solution for 72 h, with the solution replaced once, embedded in OCT compound and coronally sectioned at 30–40 μm using a Cryostat microtome (CM1950, Leica Biosystems, Germany). Sections were stored in cryoprotectant at −20 °C or processed immediately for staining.

### Immunofluorescence staining

Free-floating brain sections were blocked with 10% normal goat serum (NGS). The blocking buffer contained 0.3% Triton X-100 for c-Fos staining, but no Triton X-100 was added for Ang II or AT1R staining. Sections were then incubated overnight at 4 with the following primary antibodies, either alone or in combination: rabbit anti-c-Fos antibody (Cell Signaling Technology, 2250S, 1:500), rabbit anti-Ang II antibody (Peninsula Laboratories, T-4007, 1:500), and mouse anti-AT1R antibody (Santa Cruz, sc-515884, 1:200). After washing, sections were incubated with species-appropriate fluorescent secondary antibodies (Alexa Fluor 488/594/647, Jackson ImmunoResearch, 1:200). Sections were counterstained with 4,6-diamidino-2-phenylindole dihydrochloride (DAPI, Thermo Fisher, D1306, 1:5000) and mounted with Fluoromount-G (Southern Biotech, 0100-01).

Images were acquired using an Apotome microscope or a confocal microscope (Zeiss, Germany). Brain regions were identified according to a mouse brain atlas. c-Fos-positive cells were quantified in anatomically matched sections from the CeA, paraventricular nucleus of the hypothalamus (PVN), paraventricular nucleus of the thalamus (PVT), medial amygdala (MEA), basolateral amygdala (BLA), bed nucleus of the stria terminalis (BNST), lateral septum (LS), and locus coeruleus (LC).

Fluorescence intensity and co-localization profiles were analyzed using Image J.

### Tissue preparation and spatial RNA fluorescence in situ hybridization (RNA FISH)

Mice were deeply anesthetized with sodium pentobarbital (100 mg/kg, *i.p.*) and transcardially perfused with RNase-free PBS followed by RNase-free 4% PFA. Brains were removed, post-fixed in 4% PFA at 4 °C overnight, and then cryoprotected in 30% w/v sucrose prepared in RNase-free PBS for 72 h, with the solution replaced once. Tissues were embedded in OCT compound and coronally sectioned at 10 μm using a cryostat microtome (CM1950, Leica Biosystems, Germany) under RNase-free conditions. Sections containing the CeA were mounted onto slides and processed immediately for spatial RNA detection.

Spatial RNA detection used gene-specific probe sets targeting Slc32a1, Slc17a7, and Fos, together with DAPI nuclear counterstaining. Slc32a1 encodes vesicular inhibitory amino acid transporter and was used as a marker of GABAergic neurons. Slc17a7 encodes vesicular glutamate transporter 1 and was used as a marker of glutamatergic neurons. Fos was used as a marker of neuronal activity. The probes were designed and synthesized by Spatial FISH Biotechnology, Co., Ltd (Shenzhen, China). Before hybridization, reaction chambers were assembled on the sections to confine the reaction area and enable sequential hybridization and enzymatic reactions. The sections were dehydrated and denatured with methanol, after which hybridization buffer containing the target-specific probes was added to the chambers and incubated overnight at 37 °C. After probe hybridization, the sections were washed three times with PBST to remove unbound probes. Ligation was then performed by adding ligation mixture to the reaction chambers and incubating the sections at 25 °C for 3 h. Following ligation, the samples were washed three times with PBST and subjected to rolling circle amplification using Phi29 DNA polymerase at 30 °C overnight. Fluorescent detection probes diluted in hybridization buffer were subsequently applied to the amplified products to visualize transcript-specific signals. After fluorescent probe hybridization, the sections were dehydrated through a graded ethanol series and mounted with anti-fade mounting medium containing DAPI. Images were acquired from the CeA using a Leica THUNDER Imaging System equipped with a 20× objective lens with a numerical aperture of 0.80. Individual RNA molecules were visualized as discrete fluorescent puncta. Fluorescent signal dots were decoded according to the probe readout scheme to determine the spatial distribution of Slc32a1-, Slc17a7-, and Fos-positive cells in the CeA^[20]^.

### Stereotaxic viral injection

For all stereotaxic surgeries, adult mice were anesthetized with isoflurane anesthesia (3–4% induction, 1–2% maintenance in O_2_) and fixed in a stereotaxic apparatus (RWD, China). Body temperature was maintained at 37 °C using a heating pad throughout surgery and recovery.Viral vectors were loaded into a 10 μL Hamilton syringe connected to a 33G needle.

Injections were delivered at 50 nL/min using a microinjector pump (UMP3/Micro4, World Precision Instruments, USA), and the needle was left in place for 10 min before withdrawal. Stereotaxic coordinates were determined according to a mouse brain atlas and are reported relative to bregma.

CeA-targeted injections were performed at AP −1.55 mm, ML ±2.77 mm and DV −4.57 mm. For CeA calcium imaging, rAAV2/9-rGAD67-GCaMP6m (1.5–2.5 E+12 PFU/mL) was injected at 180 nL per site. For chemogenetic inhibition of CeA GABAergic neurons, rAAV2/9-mVGAT1-Cre (3-5 E+12 PFU/mL) was mixed 1:1 with rAAV2/9-hSyn-DIO-hM4D(Gi)-mCherry (3-5 E+12 PFU/mL) and injected bilaterally at 180 nL per side; control mice received rAAV2/9-mVGAT1-Cre (3-5 E+12 PFU/mL) mixed with rAAV2/9-hSyn-DIO-mCherry (3-5 E+12 PFU/mL). For CeA-specific *Agtr1a* knockdown, pscAAV2/9-hSyn-EGFP-miR30shRNA(*Agtr1a*)-tWPA or pscAAV2/9-hSyn-EGFP-miR30shRNA(NC)-tWPA (1.5–2.5 E+12 PFU/mL) was injected bilaterally at 180 nL per side. For calcium imaging after CeA-restricted Agtr1a knockdown, AAV-EF1a-DIO-NES-jRGECO1a was mixed 1:1 with pscAAV2/9-hSyn-EGFP-miR30shRNA(Agtr1a)-tWPA and bilaterally injected into the CeA of Gad2-Cre mice at 180 nL per side, allowing calcium responses to be monitored in CeA GABAergic neurons while Agtr1a expression was locally reduced. Control mice received AAV-EF1a-DIO-NES-jRGECO1a mixed 1:1 with the non-targeting control shRNA virus.

PVN-targeted injections were performed at AP −0.80 mm, ML ±0.20 mm and DV −4.85 mm for calcium imaging and chronic activation of glutamatergic neurons. For PVN calcium imaging, rAAV2/9-hSyn-jGCaMP8m (1.5–2.5 E+12 PFU/mL) was injected at 80 nL per site. For chronic activation of PVN glutamatergic neurons, rAAV2/9-mVGlut2-Cre (3–5 E+12 PFU/mL) was mixed 1:1 with either rAAV2/9-EF1a-DIO-mNaChBac-P2A-EGFP (3–5 E+12 PFU/mL) or rAAV2/9-EF1a-DIO-EGFP (3–5 E+12 PFU/mL) and injected bilaterally at 80 nL per side.

AAVs were allowed to express for at least 21 days before subsequent experiments, except for the pscAAV used for *Agtr1a* knockdown, which was allowed to express for at least 7 days. For *Agtr1a* knockdown experiments, viral injection was performed before CKD induction to allow stable reduction of *Agtr1a* expression during disease development. For chronic activation of PVN glutamatergic neurons, mice were first subjected to the 0.25% adenine-induced renal injury protocol for 1 week, followed by bilateral viral injection into the PVN. Samples were collected after viral expression for 3 months. Viral expression and injection sites were verified histologically after experiments, and animals with off-target expression or inaccurate targeting were excluded from analysis.

### Optical fiber implantation, fiber photometry recording and analysis

For calcium imaging experiments, an optical fiber was implanted above the viral expression site immediately after viral injection and fixed to the skull with dental cement (200 μm diameter, 0.39-NA fiber with 1.25-mm ceramic ferrule, RWD). The fiber tip was positioned approximately 0.10–0.15 mm above the injection site. Fiber placement was verified histologically after recording, and mice with misplaced fibers were excluded from analysis.

Fiber photometry was performed in freely moving mice after sufficient viral expression. Before recording, mice were habituated to handling and patch-cord connection for 3 days. Fluorescence signals were recorded using a multi-wavelength fiber-photometry system (R820, RWD, China). Fluorescence signals were acquired through the 470- and 560-nm channels to record calcium-dependent GCaMP and jRGECO1a fluorescence, respectively, while the 410-nm channel served as an isosbestic reference for correction of motion-related artifacts. Light power was maintained at 40–60 µW to reduce photobleaching.

During recording, mice were connected to the patch cord and allowed to move freely in the recording chamber. After a 5-min habituation period, baseline fluorescence was recorded for 5 min. Acute stress was then induced by a 20-s tail suspension challenge, after which mice were returned to the chamber for continued recording. Trials with signal dropout, unstable baseline or obvious movement artifacts were excluded before analysis. ΔF/F was calculated as(F − F_0_ )/F_0_ , where F_0_ was derived from a stable pre-stimulus baseline. Peak response and area under the curve (AUC) during the stress-response window were used to quantify neuronal activity.

### Ang II fluorescent tracing

To assess the CeA-associated distribution of peripherally administered Ang II under CKD conditions, mice received tail-vein injection of FAM-labeled Ang II (MedChemExpress, HY-13948F1, USA) at 1.44 mg/kg. Control groups included healthy mice receiving FAM-labeled Ang II and CKD mice receiving free 5-FAM (MedChemExpress, HY-66022, USA). Two hours after injection, mice were perfused and brains were collected for cryosectioning and fluorescence imaging. CeA fluorescence-positive area was quantified in anatomically matched sections.

### Pharmacological AT1R blockade

For systemic AT1R blockade, CKD-SUS mice received candesartan (MedChemExpress, HY-B0205) at 1 mg/kg/day by intraperitoneal (*i.p.*)injection as previously described^[21]^. Treatment began on the first day of adenine feeding and continued throughout SUS exposure. For intracerebroventricular (*i.c.v*) delivery, a guide cannula was implanted into the lateral ventricle under stereotaxic guidance (AP +1.00 mm, ML −0.50 mm and DV −2.00 mm). Candesartan was infused daily at 1 μg in 0.5 μL per mouse at 100 nL/min. The injector was left in place for 5 min after infusion. Mice with misplaced cannulas or cannula loss were excluded.

Candesartan was first dissolved in a vehicle containing 10% DMSO, 40% PEG300, 5% Tween-80 and 45% saline to obtain a clear stock solution according to product guidelines, and was then diluted with sterile saline to the required working concentration before use. Vehicle-treated mice received the same solvent mixture without candesartan.

### Multiphoton in vivo imaging

Renal intravital multiphoton imaging was performed as previously described with minor modifications^[22]^. Mice were anesthetized with isoflurane (3–4% induction, 1–2% maintenance in O_2_), and body temperature was maintained during imaging. A catheter was inserted into the jugular vein for fluorescent tracer administration. The kidney was gently exteriorized and stabilized with an intravital organ holder (OF-2000, FluoCa Scientific) to reduce respiratory and pulsatile motion. Images of superficial glomeruli were acquired using an Olympus FVMPE-RS multiphoton microscope.

To demonstrate the feasibility of renal intravital multiphoton imaging, a representative supplementary video was acquired following jugular vein injection of 5-FAM, allowing visualization of renal microvascular and nephron structures in vivo (Supplementary Video 1). Cy7-conjugated bovine serum albumin (Cy7-BSA; YS-P17010, Chongqing Yusi Pharmaceutical Technology Co., Ltd., Chongqing, China) was administered through the jugular vein catheter at a dose of 100 μL of a 5 mg/mL solution to label the plasma compartment and was excited at 1045 nm. FITC-conjugated bovine serum albumin (FITC-BSA; YS-FI012405, Chongqing Yusi Pharmaceutical Technology Co., Ltd.) was administered through the same catheter at a dose of 50 μL of a 5 mg/mL solution to assess glomerular albumin permeability and was excited at 920 nm. Following FITC-BSA administration, images of individual glomeruli were acquired at a fixed time point of 5 min after injection in mNaChBac-expressing and EGFP control-virus mice. For each animal, paired regions of interest were drawn within the glomerular capillary tuft and Bowman’s space, and mean fluorescence intensity was quantified using Image J. To account for inter-animal differences in tracer delivery and circulating fluorescence intensity, FITC-BSA fluorescence intensity in Bowman’s space was normalized to that within the corresponding glomerular capillary tuft. The resulting Bowman’s space-to-glomerular capillary tuft fluorescence intensity ratio ×100% was compared between the mNaChBac and EGFP control groups, with a higher ratio indicating greater glomerular albumin leakage.

### Quantification and statistical analysis

Statistical analyses were performed using GraphPad Prism 8.0. Data collection and analysis were performed under blinded conditions whenever possible. The value of *n* represents the number of animals unless otherwise specified. Data are presented as *mean ± SEM* unless otherwise stated.

Two-group comparisons were performed using *unpaired two-tailed Student’s t-test* or *Mann–Whitney U test* when appropriate. For comparisons among three or more groups, *one-way analysis of variance* followed by *Bonferroni post hoc tests* was used. For time-course data, *two-way repeated-measures analysis of variance* was used when both group and time were included as factors. Categorical variables were compared using *Pearson’s chi-square test*. Statistical significance was set at *p* < 0.05.

## Results

1. **Establishment and selection of the adenine-induced CKD model** To establish a clinically relevant model of CKD-associated affective comorbidity, we systematically compared three widely used CKD induction protocols: adenine diet, renal IRI, and 5/6 nephrectomy (Fig. 1A). All three models induced renal injury, although the degree of renal dysfunction varied across models. Compared with their respective controls, serum creatinine was approximately eightfold higher in adenine-fed mice and 80% higher after 5/6 nephrectomy, while remaining comparable between IRI and sham mice; similarly, BUN was approximately tenfold and twofold higher in the adenine and 5/6 nephrectomy groups, respectively, but unchanged after IRI. All three models exhibited histopathological abnormalities, including tubular atrophy, interstitial fibrosis, and glomerular lesions (Fig. 1B–F, Fig. S1 A). Notably, adenine diet-fed mice exhibited the most pronounced degree of renal dysfunction among the three groups. Beyond renal parameters, adenine diet-fed mice also displayed suggestive anxiety-like behavior in the LDB test and disrupted circadian rhythmicity at baseline—phenotypic features that parallel the emotional disturbances and sleep-wake dysregulation commonly observed in clinical CKD populations—while IRI and 5/6 nephrectomy animals showed no consistent affective or circadian changes under identical testing conditions (Fig. 1G-H, Fig. S1 B-E). Metabolic cage monitoring showed no significant differences in food intake among groups. Water intake was unchanged in adenine-fed mice but increased in 5/6 nephrectomy mice relative to sham controls (Fig. S1 F,G). Thus, adenine-induced CKD did not cause major alterations in feeding or drinking behavior. Because the adenine-induced model produced stable and robust renal dysfunction and was suitable for subsequent behavioral and circuit-level analyses, it was selected for downstream mechanistic experiments and is hereafter referred to as the CKD model.
2. **Adenine-induced CKD reveals stress-induced anxiety-like behavior under SUS** Across the different CKD models, anxiety-like behavior was not consistently observed under baseline conditions. Consistent with the clinical heterogeneity of anxiety and depression among patients with CKD, we reasoned that renal dysfunction may increase sensitivity to mild environmental stress rather than directly producing a stable affective phenotype. To test this hypothesis, we applied a subthreshold unpredictable stress (SUS) paradigm, a mild chronic stress procedure designed to be insufficient to induce significant behavioral changes in healthy control animals (Fig. 2A). SUS exposure did not significantly alter center-zone exploration in the OFT or light-compartment exploration in the LDB test in control mice. In contrast, SUS produced marked behavioral changes in CKD mice. Relative to unstressed CKD mice, CKD-SUS mice showed 54% fewer center-zone entries and a 52% reduction in distance traveled within the center zone in the OFT, together with a 54% reduction in light-compartment entries in the LDB test. Accordingly, following the same SUS exposure, CKD-SUS mice exhibited significantly lower center-zone and light-compartment exploration than control-SUS mice, indicating increased susceptibility of CKD mice to stress-induced anxiety-like behavior (Fig. 2B,C). No significant differences in immobility time during the tail suspension test were observed among the four groups (Fig. 2D), indicating that these behavioral alterations were not accompanied by depressive-like behavior.In addition, CKD mice exhibited increased freezing during the shock-paired fear-conditioning session, whereas freezing during subsequent extinction sessions, in which the conditioned stimulus was presented without foot shock, did not differ among groups (Fig. 2E; Fig. S1H). This response pattern suggests exaggerated acute threat reactivity rather than a persistent alteration in conditioned fear expression. Together, these findings indicate that the SUS paradigm remained behaviorally subthreshold in healthy control mice but unmasked anxiety-like behavior in CKD mice, supporting the conclusion that CKD increases susceptibility to mild environmental stress rather than producing a uniform baseline anxiety-like phenotype.
3. **CeA GABAergic neurons are basally activated, stress-hyperreactive, and functionally involved in CKD-associated anxiety susceptibility** To identify brain regions that are differentially activated in CKD mice, we performed c-Fos immunofluorescence across multiple emotion- and stress-related brain regions under basal conditions, including the CeA, PVN, PVT, MEA, BLA, BNST, LS, and LC. Among the regions examined, the CeA was prioritized for mechanistic investigation based on its significant CKD-associated activation and established role in anxiety-related behavior. The PVN also exhibited increased c-Fos expression in CKD mice, a finding that will be addressed in detail in a subsequent section(Fig. 3A, B). Given that previous single-cell RNA sequencing studies have shown that the CeA is composed predominantly of GABAergic neurons^[23]^, we next performed RNA FISH to characterize the cellular identity of Fos-expressing cells in CKD mice. FISH analysis revealed that approximately 16% of Slc32a1-positive GABAergic neurons in the CeA co-expressed Fos, indicating basal activation of a subset of CeA GABAergic neurons in CKD mice (Fig. 3C). These findings identified the CeA as a prominent neural target of CKD-associated basal activation. We next examined whether this basal activation was accompanied by altered CeA neuronal dynamics using fiber photometry with AAV-delivered GCaMP targeted to CeA GABAergic neurons in freely moving mice (Fig. 3D). Upon exposure to an acute stressor, CKD mice exhibited markedly enhanced CeA calcium responses compared with control mice, characterized by increased stress-evoked calcium activity and AUC (Fig. 3E, F). Consistently, CKD mice subjected to SUS also displayed elevated CeA calcium responses compared with SUS-exposed controls, indicating that CKD-associated CeA stress hyperresponsiveness persists under a mild stress challenge. To determine whether CeA GABAergic activity functionally contributes to CKD-associated stress-induced anxiety-like behavior, we next used a CeA-targeted Cre-dependent chemogenetic inhibition strategy (Fig. 4A, B). CKD mice with hM4D(Gi)-mediated inhibition of CeA GABAergic neurons showed reduced anxiety-like behavior compared with control virus-injected mice, as indicated by significantly increased center-zone exploration in the OFT and increased light-zone exploration in the LDB test (Fig. 4C, D). Together, these findings indicate that CeA GABAergic neurons exhibit basal activation and exaggerated stress responsiveness in CKD and functionally contribute to CKD-associated stress-induced anxiety-like behavior.
4. **Peripheral Ang II engages CeA AT1R signaling in CKD-associated stress-induced anxiety-like behavior** Having established the CeA as a functionally critical region, we next sought to identify the upstream signal linking peripheral renal pathology to central CeA activation. Given that the RAS is chronically overactivated in CKD, we measured circulating Ang II levels by ELISA and found significantly elevated serum Ang II in CKD mice compared to controls (Fig. 5A). To determine whether this peripherally elevated Ang II accesses the central nervous system, we measured Ang II levels in CSF and quantified Ang II immunofluorescence intensity within the CeA. Neither CSF Ang II concentration nor CeA immunofluorescence signal differed significantly between CKD and control animals (Fig. S2A–C), suggesting that under steady-state conditions, peripheral Ang II does not stably accumulate in the central compartment at measurable levels. However, the absence of detectable static central accumulation does not exclude transient regional access. We therefore performed peripheral fluorescent Ang II tracing using FAM-labeled Ang II, in which Ang II was conjugated to a FAM fluorophore. CKD mice receiving free FAM were included to control for nonspecific fluorophore retention or accumulation. Following peripheral administration, FAM-Ang II-associated fluorescence was increased in the CeA of CKD mice compared with control mice, supporting enhanced regional enrichment of circulating Ang II-associated signal under CKD conditions (Fig. 5B,C). Immunofluorescence further revealed abundant AT1R expression within the CeA, with c-Fos-positive cells located in AT1R-enriched regions (Fig. 5D, E). Although this spatial association does not establish direct receptor-mediated neuronal activation, it supports the plausibility that locally enriched Ang II may engage CeA AT1R signaling. This observation prompted us to examine the functional contribution of local CeA AT1R signaling. To assess the contribution of local CeA Agtr1a signaling to the manifestation of the behavioral phenotype, we first performed CeA-restricted *Agtr1a* knockdown by stereotaxic injection of AAV-shRNA targeting *Agtr1a* prior to CKD induction, with scramble shRNA-injected mice serving as controls (Fig. 6A). Knockdown efficiency within the CeA was confirmed by immunofluorescence (Fig. S2D,E). Under the SUS paradigm, CKD mice with CeA-specific *Agtr1a* knockdown exhibited attenuated anxiety-like behavior in both the OFT and LDB tests (Fig. 6B,C). Calcium imaging further showed that *Agtr1a* knockdown attenuated the exaggerated stress-evoked CeA calcium responses observed in CKD mice (Fig. 6D–F), supporting a functional contribution of local CeA AT1R signaling to altered stress responsiveness and anxiety-like behavior. We next examined whether broader pharmacological AT1R blockade produced similar behavioral effects by administering candesartan either systemically or intracerebroventricularly using literature-based dosing regimens. Neither treatment significantly altered anxiety-like behavior under the conditions tested (Fig. S3A–F). Because regional drug exposure and CeA AT1R occupancy were not directly quantified, these findings should be interpreted as regimen-specific negative results rather than definitive evidence against Ang II–AT1R involvement.
5. **CKD enhances PVN stress responses, whereas sustained experimental PVN activation increases renal injury markers** In addition to the CeA abnormalities identified above, our initial c-Fos mapping revealed increased basal activation of the PVN in CKD mice (Fig. 7A). The PVN is a major autonomic and neuroendocrine output node, and previous studies have established its role in regulating renal sympathetic activity and cardiorenal homeostasis^[14]^. These observations prompted us to examine whether CKD alters PVN stress responsiveness and whether sustained activation of this central output region can influence renal injury. Fiber photometry calcium imaging during acute stress exposure showed that CKD mice exhibited enhanced stress-evoked PVN calcium responses compared with control mice, as reflected by an increased positive area under the curve (AUC) of the calcium signal (Fig. 7B–D). Together, these findings identify the PVN as an additional stress-responsive region sensitized by CKD. We next examined whether sustained PVN activation could contribute to a potential brain-to-kidney feedback process. Using a mild renal injury model induced by 0.25% adenine feeding for one week, we activated PVN glutamatergic neurons through stereotaxic delivery of an AAV encoding the bacterial sodium channel mNaChBac, thereby producing sustained cell-autonomous excitation and modeling prolonged PVN hyperactivity (Fig. 7E). Following the activation period, mNaChBac-expressing mice exhibited increased levels of the early tubular injury markers KIM-1 and NGAL, together with elevated UACR, compared with EGFP control-virus mice, whereas serum creatinine and BUN remained comparable between groups (Fig. 7F–J). Renal intravital multiphoton imaging further revealed increased fluorescence intensity within Bowman’s space in the PVN activation group, consistent with enhanced glomerular albumin leakage (Fig. 7K, L; Supplementary Video 2). This experiment was designed to test the renal consequences of experimentally sustained PVN activation rather than to reproduce the magnitude or temporal pattern of PVN activity occurring naturally in CKD. The findings provide proof-of-concept evidence that sustained activation of PVN glutamatergic neurons can exacerbate early renal injury in mice with mild pre-existing renal injury. Although the downstream autonomic pathways were not directly examined, these results support the possibility that maladaptive central stress-output activity contributes to brain-to-kidney feedback in CKD.

**Figure 1.**
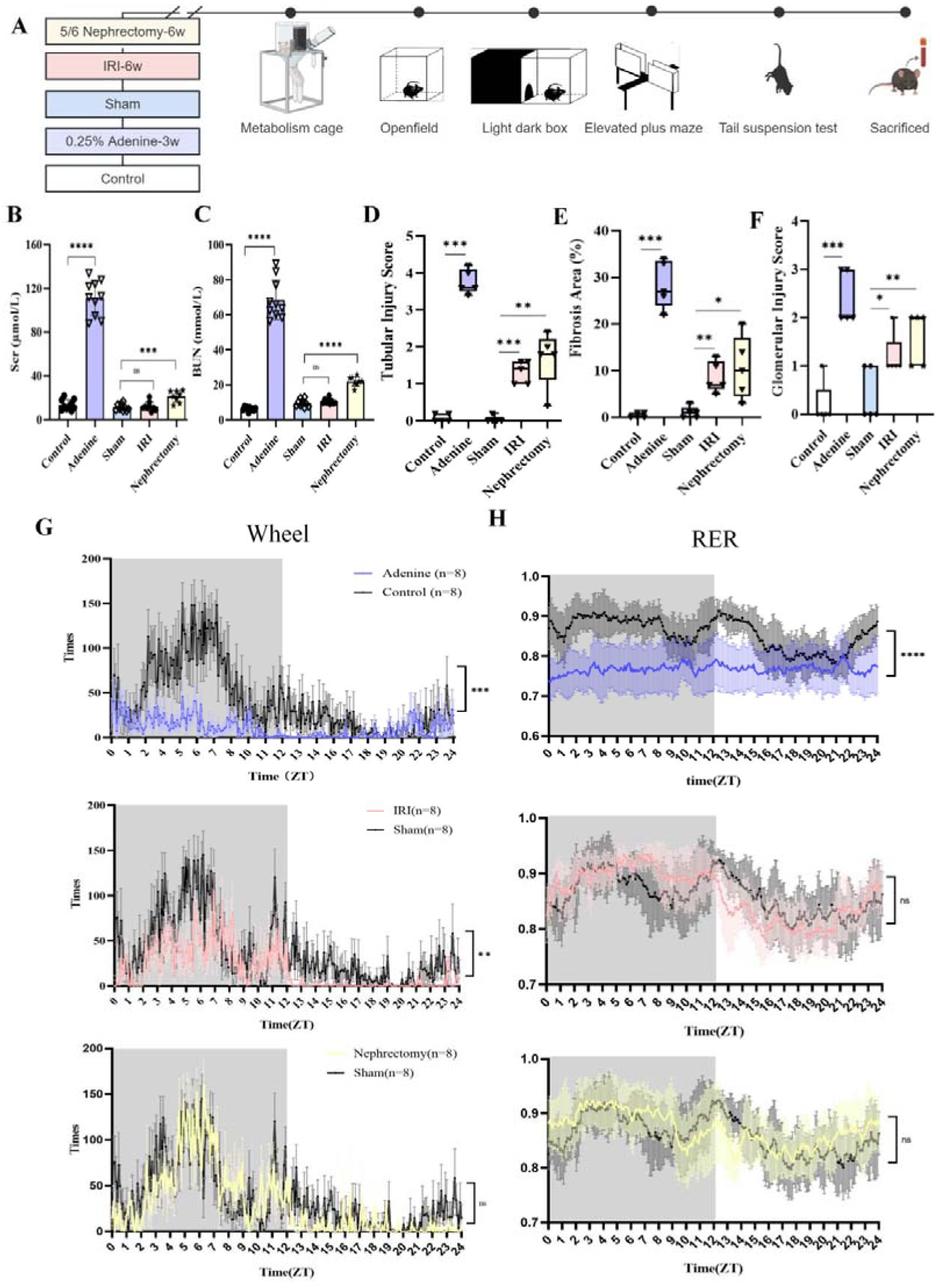
Renal function and circadian rhythmicity across three chronic kidney disease (CKD) models. (A) Experimental timeline of CKD models. Metabolic cage analysis and behavioral tests were performed 6 weeks after 5/6 nephrectomy, IRI, or sham surgery, and 3 weeks after 0.25% adenine treatment or control intervention, followed by tissue collection (B-C) Serum biochemical indices of renal function: (B) serum creatinine (Scr) and (C) blood urea nitrogen (BUN) levels across five experimental groups. n = 10 per group, except for the IRI group (n = 9) and the 5/6 nephrectomy group (n = 8). (D-F) Semi-quantitative histopathological scoring of renal tissue: (D) tubular injury score based on H&E staining; (E) percentage of interstitial fibrosis area based on Masson’s trichrome staining; (F) glomerular injury score based on PAS staining. (n = 4 per group). (G) Continuous 24-hour locomotor activity rhythms (activity counts per hour) in adenine diet-fed, IRI, and nephrectomy model mice, respectively, compared with their corresponding controls. The grey area represents the “dark phase”. (n = 8 per group).(H) Continuous 24-hour respiratory exchange ratio (RER = VCO /VO ) rhythms reflecting diurnal dynamics of metabolic substrate utilization in adenine diet-fed, IRI, and nephrectomy model mice, respectively, compared with their corresponding controls. The grey area represents the “dark phase”. (n = 8 per group).Data are presented as mean ± SD or min-to-max range (panels D-F). Statistics used two-tailed unpaired t tests for panel B-C, Mann–Whitney U test for panel D-F and two-way repeated-measures analysis of variance for panel G-H.“ns” indicates no significant difference; *p < 0.05, **p < 0.01, ***p < 0.001, ****p < 0.0001.

**Figure 2.**
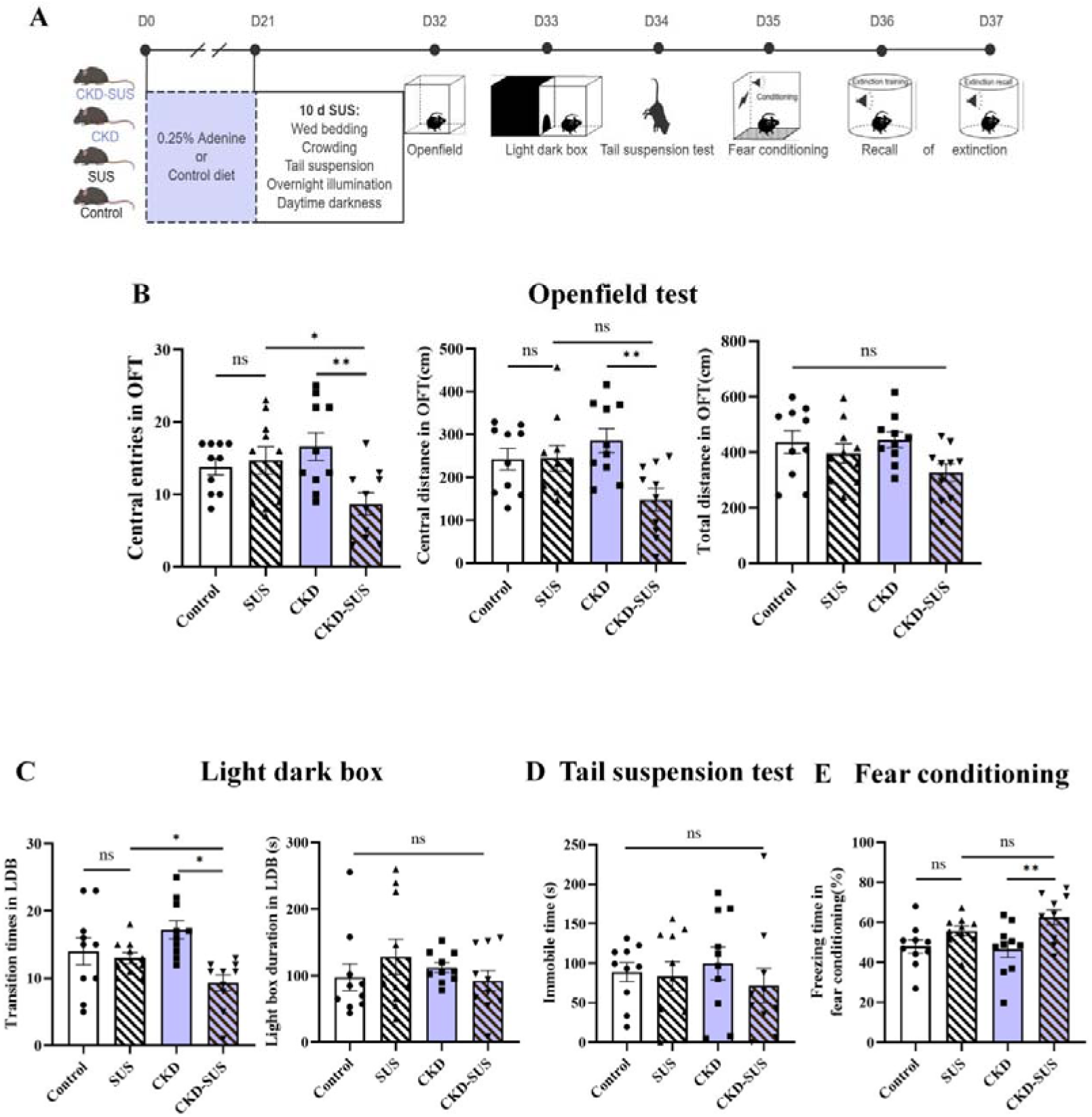
Chronic kidney disease (CKD) induces anxiety susceptibility under subthreshold unpredictable stress (SUS). (A) Experimental design of the SUS paradigm. Four groups were included (n = 10 per group). Mice were subjected to 10 days of unpredictable mild stress, followed by behavioral testing including open field test (OFT), light-dark box (LDB), tail suspension test (TST), and contextual fear conditioning. (B) OFT performance, including total distance, number of entries into and distance traveled within the center zone. (C) LDB performance, including number of entries into and time spent in the light compartment. (D) Immobility time in the TST across groups. (E) Percentage of freezing during fear conditioning acquisition. Data are presented as mean ± SEM. Statistics used one-way analysis of variance followed by Bonferroni post hoc tests for behavioral comparisons. “ns” indicates no significant difference, *p < 0.05, **p < 0.01, ***p < 0.001.

**Figure 3.**
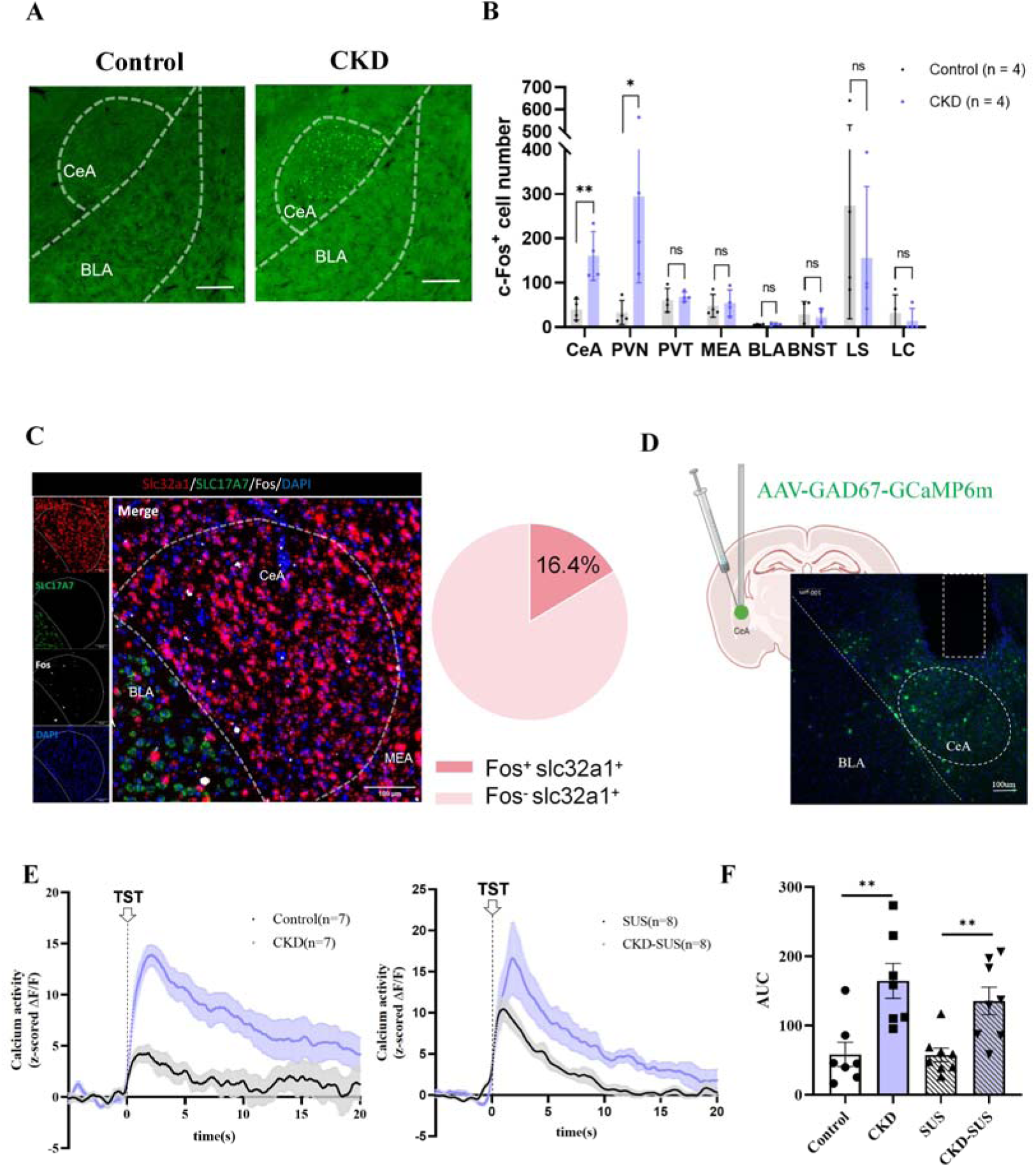
Central amygdala (CeA) GABAergic neurons are basally activated and exhibit stress hyperreactivity in chronic kidney disease (CKD) mice. (A) Representative c-Fos immunofluorescence images in the CeA from Control and CKD mice (scale bar: 200 μm).(B) Quantification of c-Fos-positive neurons across brain regions associated with stress and emotion.(C) Left, representative FISH images showing Slc32a1+ GABAergic neurons (red), Slc17a7+ glutamatergic neurons (green), Fos+ cells (white), and DAPI (blue) in the CeA of CKD mice. Right, pie chart showing the percentage of Slc32a1+ neurons co-expressing Fos (n=3). (scale bar: 100 μm).(D) Schematic of in vivo fiber photometry calcium imaging (scale bar: 100 μm).(E) Mean peri-event traces of z-scored ΔF/F aligned to the onset of acute tail-suspension stress (TST).(F) Area under the curve (AUC) of calcium signals during stress. Data are presented as mean ± SD (B) or mean ± SEM (F). Statistics used two-tailed unpaired t tests.“ns” indicates no significant difference, *p < 0.05, **p < 0.01.

**Figure 4.**
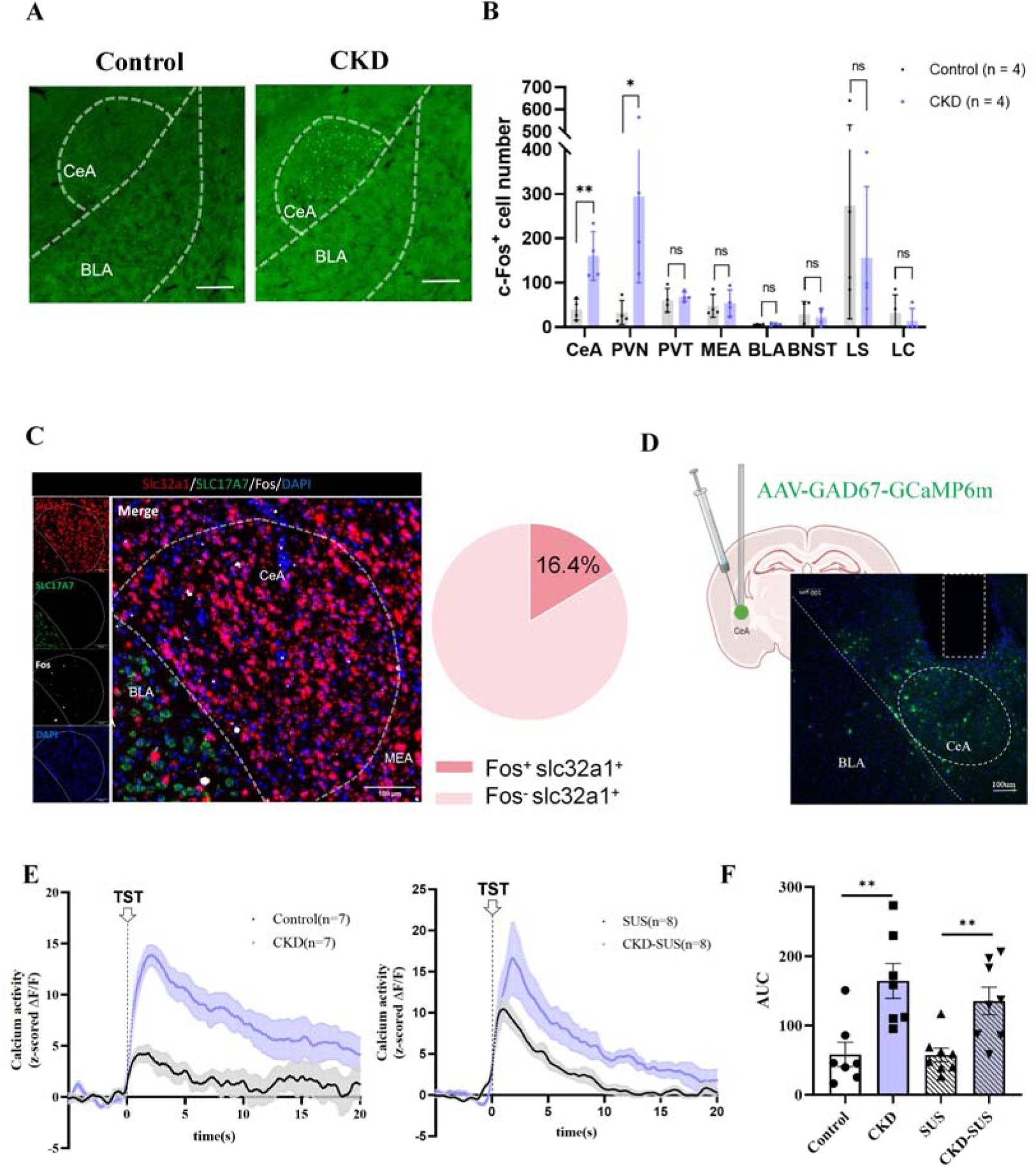
Chemogenetic inhibition of central amygdala (CeA) GABAergic neurons attenuates anxiety susceptibility in chronic kidney disease (CKD) mice. (A) Experimental timeline for chemogenetic inhibition (n = 8 per group). C57 mice received bilateral CeA AAV injections 7 days before CKD-SUS modeling; DCZ was systemically administered before behavioral testing for DREADD manipulation.(B) Viral strategy and expression. Left: injection sites; right: representative fluorescence images (scale bar: 200 μm).(C) Open field test (OFT) performance, including center-zone entries and distance.(D) Light-dark box (LDB) performance, including entries into and time spent in the light compartment.Data are presented as mean ± SEM. Statistics used two-tailed unpaired t tests.“ns” indicates no significant difference, *p < 0.05, **p < 0.01.

**Figure 5.**
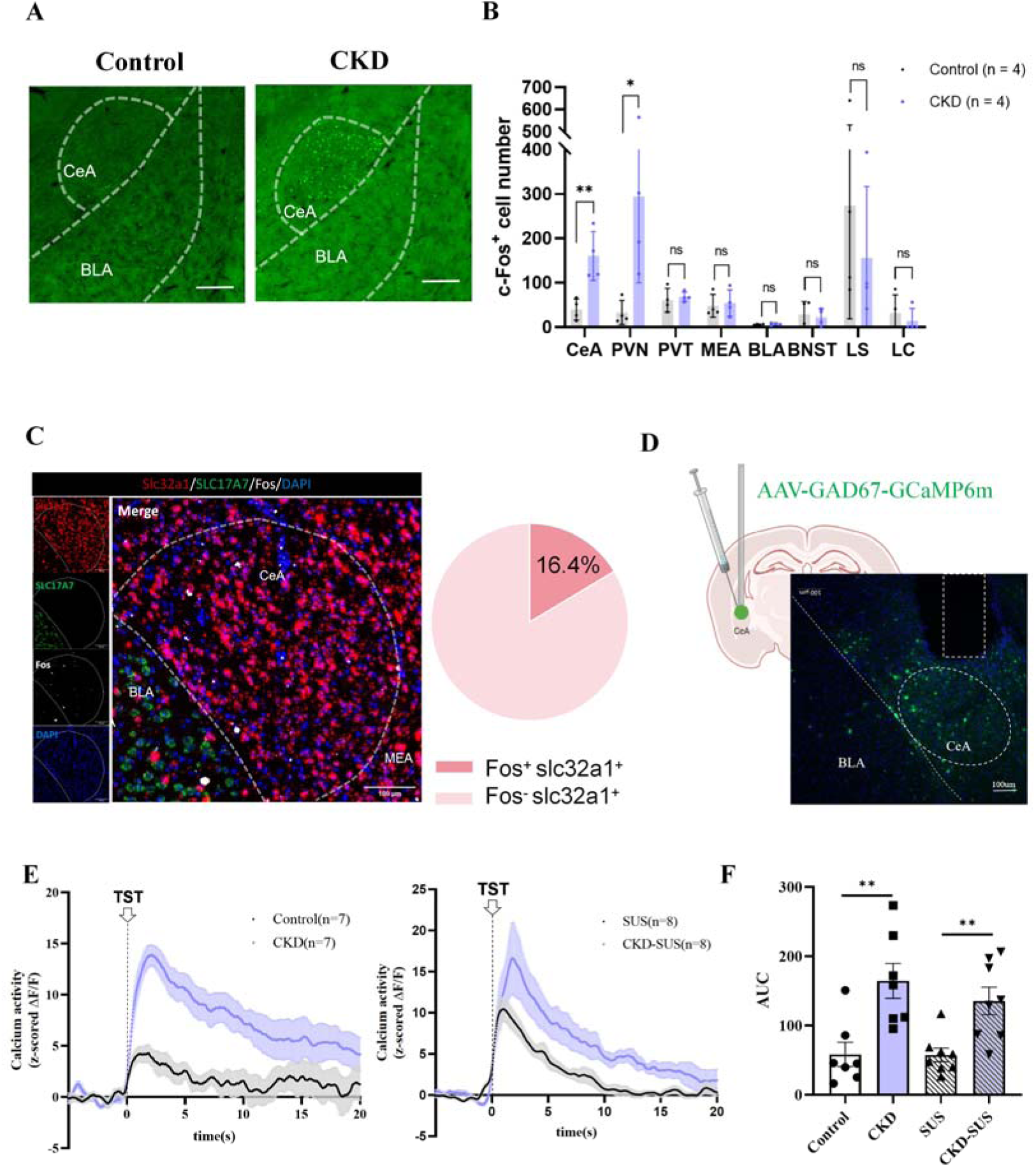
Chronic kidney disease (CKD) elevates circulating Ang II and enhances central amygdala (CeA) accumulation of intravenously administered FAM-Ang II-associated fluorescence. (A) Serum Ang II levels in Control, CKD, SUS and CKD-SUS mice (n=10 per group).(B-C) Quantification and representative images of fluorescence accumulation in the CeA (n = 5; scale bars: left 100 μm and right 50 μm).(D) Representative immunofluorescence images of AT1R (red) and c-Fos (white) expression in CeA sections, with DAPI nuclear staining (blue) (scale bar: 50 μm).(E) Representative co-localization curve corresponding to the region indicated by the green arrow (pointing) in panel (D).Data are presented as mean ± SEM. Statistics used one-way analysis of variance followed by Bonferroni post hoc tests. *p < 0.05, **p < 0.01.

**Figure 6.**
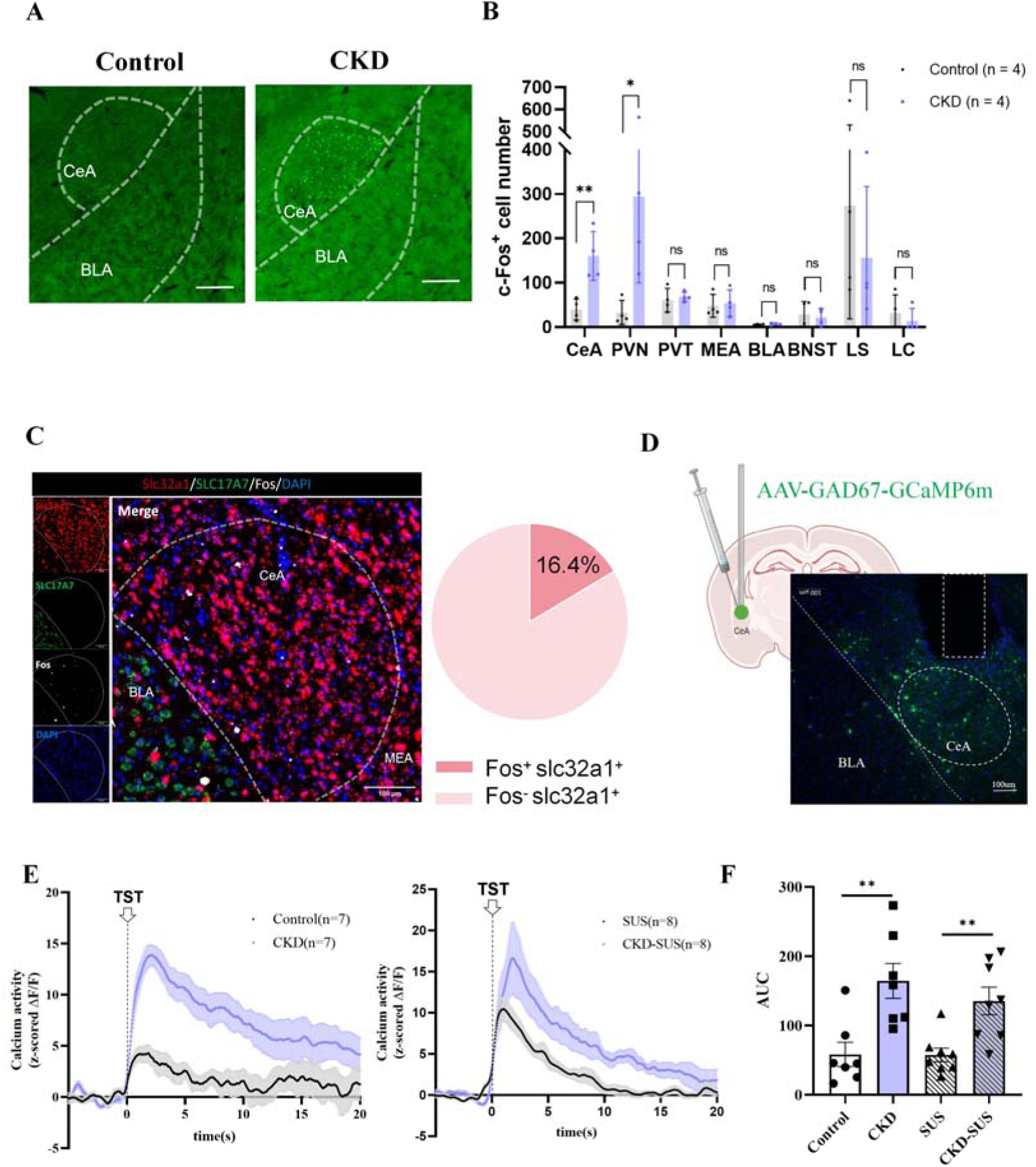
Central amygdala (CeA)-specific Agtr1a knockdown reduces anxiety susceptibility in chronic kidney disease (CKD) mice. (A) Experimental design. Wild-type mice received bilateral CeA injection of AAV carrying Agtr1a shRNA or control vector prior to CKD-SUS induction, followed by behavioral testing (n = 12 per group).(B) Open field test (OFT) performance, including center-zone entries and distance. (C) Light-dark box (LDB) performance, including entries into and time spent in the light compartment.(D) Experimental design. Schematic of in vivo fiber photometry calcium imaging after bilateral CeA Agtr1a knockdown in CKD GAD2-cre mice. (scale bar: 100 μm).(E) Mean peri-event traces of z-scored ΔF/F aligned to the onset of acute tail-suspension stress (TST) in NC and Agtr1a-knockdown mice. NC served as the control group.(F) Area under the curve (AUC) of calcium signals during stress.Data are presented as mean ± SEM. Statistics used two-tailed unpaired t tests. *p < 0.05, **p < 0.01, ***p < 0.001.

**Figure 7.**
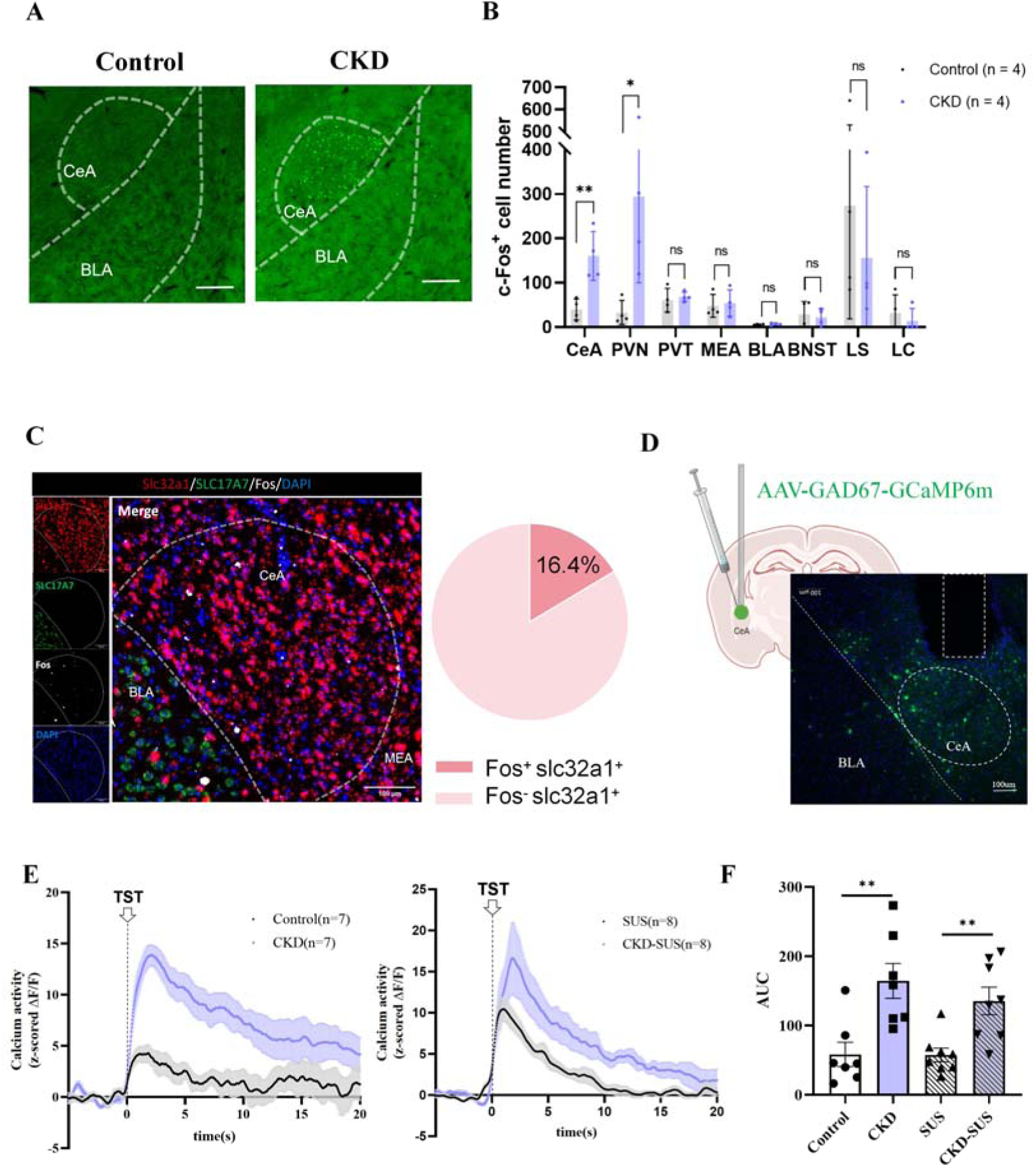
Chronic kidney disease (CKD) remodels paraventricular nucleus (PVN) stress responsiveness, and sustained PVN activation exacerbates early renal injury. (A) Representative c-Fos immunofluorescence images in the PVN from Control and CKD mice (scale bar: 200 μm).(B) Schematic of fiber photometry recording in the PVN, with representative viral expression and fiber placement (scale bar: 100 μm).(C) Mean peri-event traces of z-scored ΔF/F aligned to the onset of acute tail-suspension stress (TST).(D) Area under the curve (AUC) of positive stress-evoked calcium responses above baseline. (E) Experimental design for chronic activation of PVN neurons using AAV-mediated mNaChBac expression. Left: injection sites; right: representative fluorescence images (scale bar: 200 μm). (F–J) Biochemical and injury markers following chronic PVN activation, including Scr, BUN, KIM-1, NGAL, and UACR (n=8).(K) Representative intravital multiphoton microscopy images of the kidney in the Control and NachBac groups. Glomeruli are indicated by white dashed circles and white arrows, while Bowman’s capsule is indicated by green arrows. In the NachBac group, green fluorescent albumin signal was observed within Bowman’s space, indicating albumin leakage across the glomerular filtration barrier. In contrast, little or no albumin leakage was detected in the Control group.(L) Bowman’ s space-to-glomerular capillary tuft FITC-BSA fluorescence intensity ratio ×100% at 5 min after intravenous FITC-BSA administration in the EGFP control and mNaChBac groups. (n=4).Data are presented as mean ± SEM. Statistics used two-tailed unpaired t tests. “ns” indicates no significant difference; *p < 0.05, **p < 0.01, ***p < 0.001.

## Discussion

Here, we show that CKD promotes susceptibility to mild stress, leading to the emergence of anxiety-like behavior through a CeA-centered mechanism involving local Ang II–AT1R signaling. CKD did not impose a uniform anxiety-like phenotype at baseline; instead, it lowered the threshold at which mild stress triggered maladaptive anxiety-like behavior. The CeA emerged as a disease-sensitized affective node, with enhanced basal activation and exaggerated stress-evoked calcium responses in CeA GABAergic neurons. The data further implicate peripheral Ang II and local CeA AT1R signaling in this phenotype. CKD increased circulating Ang II and enhanced the CeA accumulation of peripherally administered FAM-Ang II-associated signal, suggesting increased regional access of circulating Ang II-related signals under disease conditions. Moreover, CeA-restricted Agtr1a knockdown attenuated both anxiety-like behavior and exaggerated stress-evoked CeA calcium responses. Nevertheless, the precise route by which circulating Ang II-related signals access and engage the CeA remains to be defined. In parallel, CKD remodeled PVN stress responsiveness, and sustained PVN glutamatergic activation aggravated early renal injury, suggesting that central stress-output circuits may feed back onto the injured kidney. Together, these findings provide exploratory evidence consistent with a potential brain-to-kidney feedback component in which renal dysfunction sensitizes central affective circuits, while maladaptive central output may further amplify renal injury.

### CKD Lowers the Threshold for Stress-Induced Anxiety-like Behavior

The behavioral data suggest that CKD functions as a stress-sensitizing condition rather than a standalone anxiety model. In the initial model comparison, adenine-induced CKD produced the most robust renal dysfunction and histopathological injury among the models tested, accompanied by disruption of circadian and metabolic rhythms. However, baseline anxiety-like behavior was not consistently expressed across assays, even in this model. Similar inconsistencies have been reported in subtotal nephrectomy models, where baseline affective phenotypes appear to vary across studies^[24,25]^. In adenine-induced CKD, a mixed affective profile has also been described, including depression-like behavior alongside an apparently anxiolytic pattern in the LDB test, as reflected by increased transitions and greater time spent in the light compartment^[26]^.

This finding is important because it indicates that CKD alone may not be sufficient to generate stable affective abnormalities under resting laboratory conditions. The subthreshold unpredictable stress paradigm revealed this stress-dependent phenotype: stressors that were behaviorally ineffective in healthy mice induced reduced exploration in anxiogenic environments and enhanced acute freezing responses in CKD mice. This pattern suggests reduced affective resilience (i.e., increased vulnerability to otherwise manageable stressors), in which otherwise tolerable stressors evoke anxiety-like avoidance. This concept parallels evidence from chronic disease populations showing that diminished psychological resilience is associated with greater depression, anxiety, and stress-related affective burden^[27,28]^. Together, these findings support a threshold-shift model of CKD-associated anxiety-like behavior.

### CeA GABAergic Circuits Provide a Substrate for CKD-Induced Anxiety-like Vulnerability

The CeA provides a plausible circuit substrate for this loss of affective resilience. Inhibitory microcircuits within the amygdala, particularly within the CeA, are central to threat processing, fear expression and anxiety-related behavioral output^[29,30]^. Our data extend this framework by showing that a peripheral renal disease state can sensitize CeA GABAergic stress responses. In CKD mice, c-Fos mapping identified enhanced CeA activation, and co-localization analyses support recruitment of CeA GABAergic neurons. Fiber photometry further showed that CeA GABAergic neurons displayed exaggerated calcium responses to acute stress, indicating that CKD increases stress-evoked neuronal gain in this region. Importantly, chemogenetic inhibition of CeA GABAergic neurons attenuated anxiety-like behavior in CKD-SUS mice, supporting a functional requirement for this population in the behavioral phenotype.

Because our viral strategy targeted broad CeA GABAergic populations, the specific CeA subnuclei and molecularly defined neuronal subtypes responsible for this phenotype remain unresolved. These findings suggest that CKD shifts the CeA from a calibrated threat-processing structure into a disease-sensitized gain-control node that converts mild stress input into exaggerated anxiety-like behavioral output. This raised the question of which kidney-derived signal drives CeA sensitization in CKD.

### CeA Ang II–AT1R Signaling Contributes to CKD-Associated Stress Susceptibility

Brain Ang II–AT1R signaling has been linked to stress responsiveness, HPA-axis activation, sympathetic regulation and anxiety-like behavior in previous studies^[31]^. CKD has also been associated with impaired blood–brain barrier integrity, which may create a permissive condition for circulating Ang II-related signals to influence vulnerable brain regions^[6]^. Our findings extend this framework by suggesting that Ang II-related signaling in CKD may be spatially concentrated within the CeA rather than reflected by sustained global accumulation in the central compartment.

CKD increased circulating Ang II, but steady-state measurements of CSF Ang II and static CeA Ang II immunofluorescence did not show robust elevation. In contrast, peripheral administration of fluorescently labeled Ang II revealed enhanced CeA signal in CKD mice, supporting disease-dependent access or enrichment of circulating Ang II within this limbic region. Because BBB permeability, vascular leakage, and cellular transport routes were not directly examined, these findings do not define the precise entry mechanism. Nevertheless, the localization of c-Fos-positive cells within AT1R-enriched CeA regions supports the plausibility that peripherally derived Ang II-related signals may engage local AT1R signaling. CeA-restricted *Agtr1a* knockdown attenuated anxiety-like behavior and exaggerated stress-evoked CeA calcium responses in CKD mice. These spatially targeted findings support a contribution of local CeA AT1R signaling to altered CeA stress responsiveness and the associated behavioral phenotype.

Previous work has proposed that AT1R blockers may reduce excessive stress responses, neuroinflammation or anxiety-related behaviors in some models ^[31,32]^. However, behavioral effects of systemic AT1R blockade treatment are not uniform across disease contexts, with some studies reporting limited or absent benefit in affective phenotypes ^[33]^. In our CKD-SUS model, the lack of behavioral rescue after systemic or ICV candesartan suggests that broad AT1R blockade, at least under the dosing and timing conditions used here, is insufficient to reverse CKD-associated stress-induced anxiety-like behavior. These negative pharmacological findings should be interpreted cautiously, because the present study did not directly assess blood pressure responses, renal biochemical effects, regional candesartan distribution, or CeA receptor occupancy in these cohorts.

### PVN stress-output sensitization promotes renal injury

Beyond the CeA, our data suggest that CKD also alters hypothalamic stress-output circuits. The PVN is a major autonomic and neuroendocrine control center, and prior work has implicated PVN-related pathways in the regulation of renal sympathetic output and cardiorenal homeostasis ^[14,34]^.

In our study, CKD increased basal PVN activation and enhanced stress-evoked PVN calcium responses, consistent with central stress-output sensitization.We focused on PVN glutamatergic neurons because Vglut2 marks a broad excitatory PVN population that includes several neuropeptide-defined stress- and arousal-related subpopulations, including CRH-, OXT-, and PDYN-expressing neurons^[35]^. To determine whether sustained activation of this output node is sufficient to influence renal pathology, we chronically activated PVN glutamatergic neurons and found evidence of aggravated renal injury despite relatively preserved conventional measures of renal function. Consistent with the increase in urinary albumin indices, intravital renal multi-photon imaging revealed greater FITC-albumin signal within Bowman’s space in PVN-activated mice, supporting enhanced glomerular albumin leakage. Sympathetic renal efferents represent a plausible downstream route by which stress- or emotion-related PVN overactivation may aggravate renal injury. Future studies using direct nerve recording, renal denervation, or adrenergic blockade will be required to test whether renal sympathetic output mediates this brain–kidney effect. These findings provide proof-of-concept evidence that maladaptive central stress output can feed back onto the kidney.

In conclusion, our study identifies a localized Ang II–CeA AT1R mechanism through which CKD primes stress-induced anxiety susceptibility. This pathway links peripheral renal dysfunction to CeA GABAergic stress hyperresponsiveness and explains how mild environmental stress can precipitate anxiety-like behavior in the CKD state. By further showing that sustained PVN activation aggravates early renal injury and is associated with increased glomerular albumin leakage, our findings provide exploratory evidence consistent with a potential brain-to-kidney feedback component. This framework suggests that CKD-associated neuropsychiatric comorbidity may arise from disease-sensitized brain circuits and points to region-specific circuit mechanisms that may be missed by systemic RAS blockade.

## Acknowledgement

Not applicable.

## Author contribution

YL and YH contributed equally to this work. YL and YH designed and performed the major experiments, analyzed the data, prepared the figures, and drafted the manuscript. ZW and XZ assisted with animal experiments, behavioral assessments, tissue collection, and data analysis. LZ and MH contributed to experimental design, technical support, and interpretation of the data. NH and FY provided experimental resources, supervised the study, and contributed to the conceptual development of the project. LZ, NH, HM, and FY critically revised the manuscript. All authors reviewed and approved the final version of the manuscript.

## Ethics approval

All procedures were carried out in accordance with the protocols approved by the Ethics Committee of the Animal Care and Use Committee of Shenzhen People’s Hospital (AUP-240417-LYT-256-01) and Shenzhen Institutes of Advanced Technology, Chinese Academy of Sciences (SIAT-IACUC-190219-NS-YF-A0582).

## Consent for publication

Not applicable.

## Data availability

The raw data supporting the conclusions of this article will be made available by the authors on request.

---

**Supplementary Figure 1.**
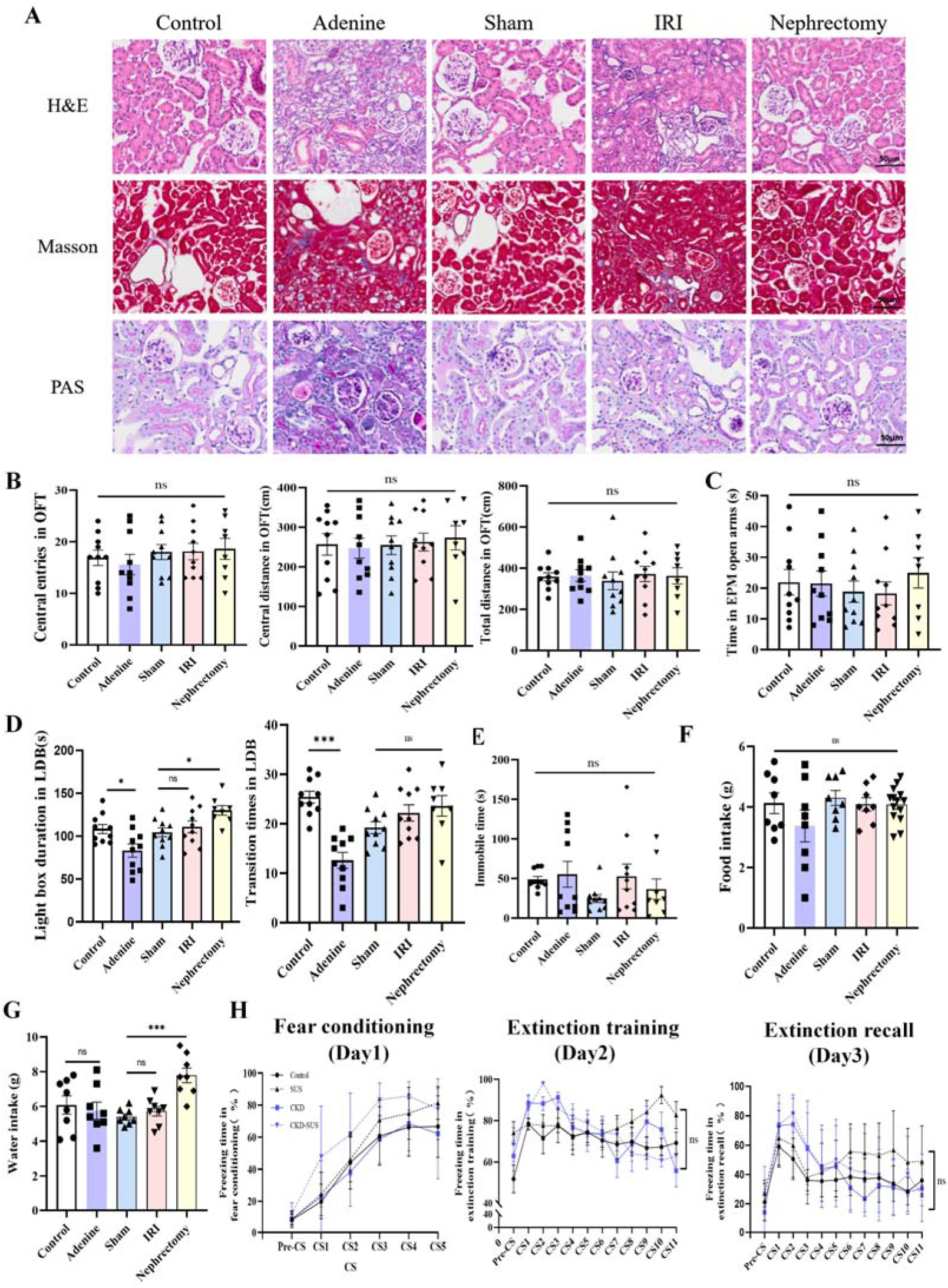
Histopathological and behavioral characterization of chronic kidney disease(CKD) models. (A) Representative histological images of renal tissue stained with H&E (top), Masson’s trichrome (middle), and PAS (bottom) from Control, Adenine, Sham, IRI, and Nephrectomy groups. Scale bar: 50 μm.(B) Open field test (OFT): total distance, number of entries into and distance traveled within the center zone across groups.(C) Elevated plus maze (EPM): time spent in the open arms across groups.(D) Light-dark box (LDB) : including number of entries into and time spent in the light compartment across groups. (E) Tail suspension test (TST): total immobility time across groups.(F) Food and (G) water intake during metabolic cage monitoring across CKD models (n=8).(H) Percentage of freezing during each conditioned stimulus (CS) presentation in the fear test. Data are presented as mean ± SEM. Statistics used one-way analysis of variance followed by Bonferroni post hoc tests and two-way repeated-measures ANOVA for panel H.“ns” indicates no significant difference, *p < 0.05, **p < 0.01.

**Supplementary Figure 2.**
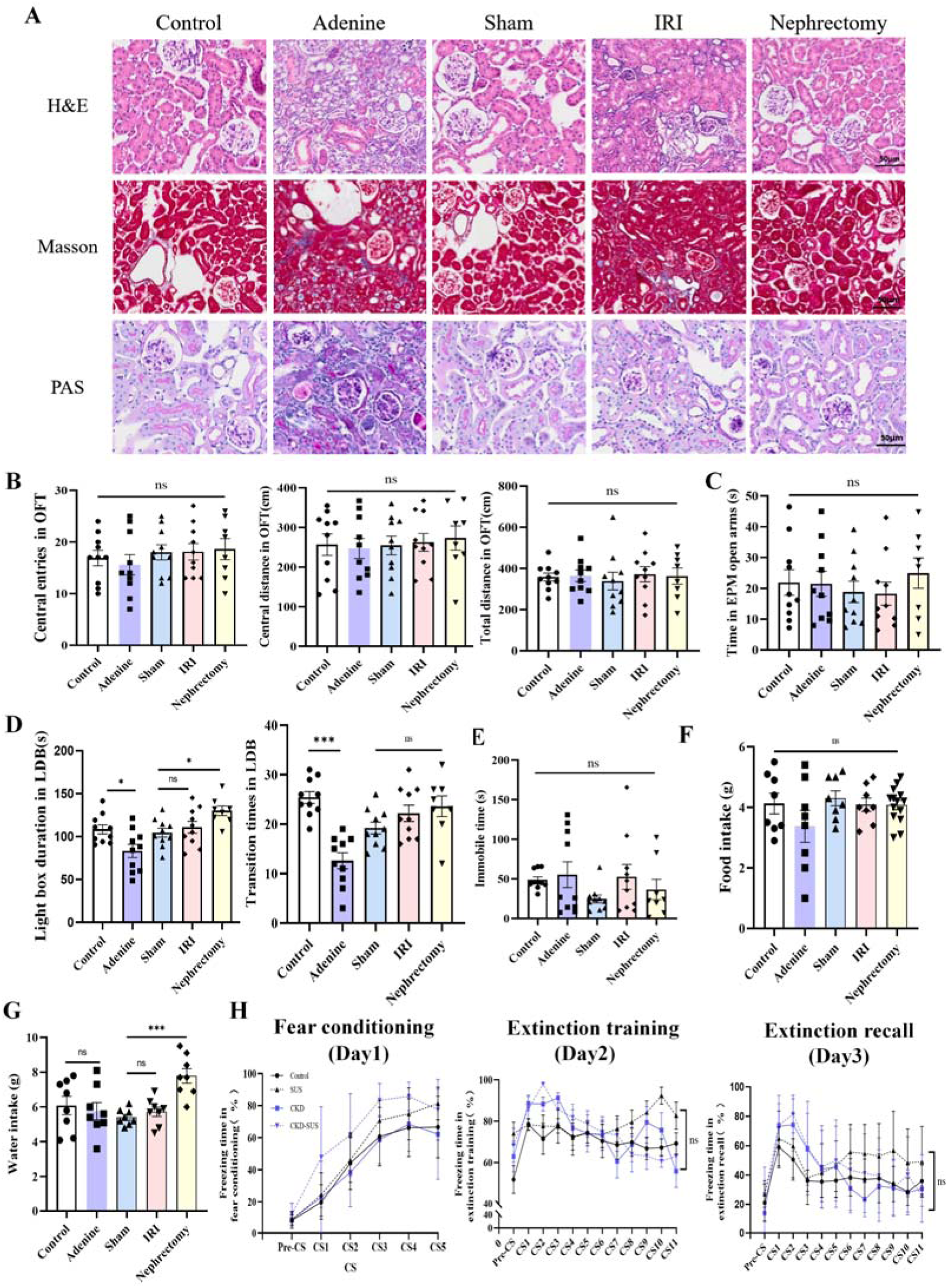
Central Ang II measurements and validation of CeA Agtr1a knockdown. (A) Cerebrospinal fluid (CSF) Ang II levels. Due to limited CSF volume, each data point represents pooled samples from 3–5 mice (detection limit: 0.1 pg/mL).(B) Representative immunofluorescence images of Ang II in the central amygdala (CeA) (scale bars: 100 μm and 50 μm).(C) Quantification of Ang II fluorescence-positive area in the CeA (n = 4).(D) Representative images of viral transduction and AT1R expression in the CeA (n = 3). Green: virus; red: AT1R; blue: DAPI (scale bar: 100 μm).(E) Quantification of AT1R expression in the CeA after viral transduction (n = 3).Data are presented as mean ± SEM. Statistics used two-tailed unpaired t tests.“ns” indicates no significant difference, **p < 0.01.

**Supplementary Figure 3.**
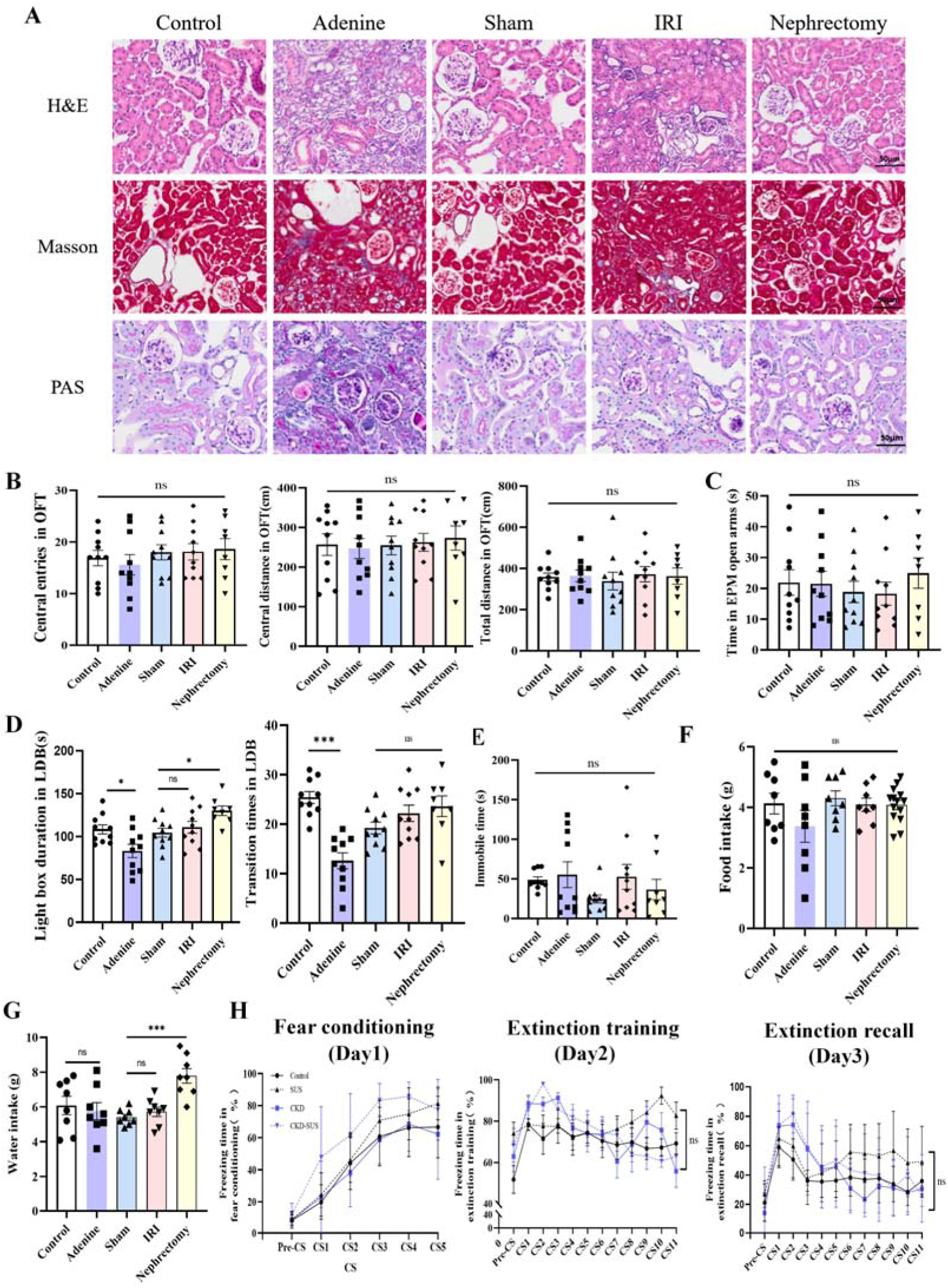
Systemic and central angiotensin receptor blocker (ARB) administration in chronic kidney disease (CKD) mice. (A) Experimental design for systemic (intraperitoneal) ARB administration (n=10).(B) Open field test (OFT) performance, including center-zone entries and distance following systemic treatment. (C) Light-dark box (LDB) performance, including entries into and time spent in the light compartment following systemic treatment. (D) Experimental design for intracerebroventricular (ICV) ARB administration (n=10).(E) OFT performance following ICV treatment. (F) LDB performance following ICV treatment.Data are presented as mean ± SEM. Statistics used two-tailed unpaired t tests.“ns” indicates no significant difference.

**Supplementary Video 1. Intravital multiphoton imaging of the kidney after vein injection of 5-FAM.** Representative intravital multiphoton microscopy video demonstrating the feasibility of the renal two-photon imaging approach established in this study. Following jugular vein injection of the green fluorescent tracer 5-FAM, renal microvascular and nephron structures were clearly visualized, including the afferent arteriole, glomerulus, efferent arteriole, Bowman’s capsule, and renal tubules. The dynamic fluorescence imaging confirms successful in vivo labeling and enables real-time observation of renal glomerular and tubular structures.

**Supplementary Video 2. Intravital multiphoton imaging of albumin leakage in the NachBac group.** Representative intravital multiphoton microscopy video showing green fluorescent albumin leakage within Bowman’s space in the NachBac group. The white arrow indicates Bowman’s capsule, where accumulation of green fluorescence was observed, suggesting increased albumin permeability across the glomerular filtration barrier.

## Reference

[1] Kovesdy C P. Epidemiology of chronic kidney disease: an update 2022[J]. Kidney International Supplements, 2022, 12(1): 7–11.

[2] Adejumo O A, Edeki I R, Sunday Oyedepo D, et al. Global prevalence of depression in chronic kidney disease: a systematic review and meta-analysis[J]. Journal of Nephrology, 2024, 37(9): 2455–2472.

[3] Loosman W L, Rottier M A, Honig A, et al. Association of depressive and anxiety symptoms with adverse events in Dutch chronic kidney disease patients: a prospective cohort study[J]. BMC Nephrology, 2015, 16(1): 155.

[4] Derrett S, Samaranayaka A, Schollum J B W, et al. Predictors of Health Deterioration Among Older Adults After 12 Months of Dialysis Therapy: A Longitudinal Cohort Study From New Zealand[J]. American Journal of Kidney Diseases, 2017, 70(6): 798–806.

[5] Adesso S, Paterniti I, Cuzzocrea S, et al. AST-120 Reduces Neuroinflammation Induced by Indoxyl Sulfate in Glial Cells[J]. Journal of Clinical Medicine, 2018, 7(10): 365.

[6] Bobot M, Thomas L, Moyon A, et al. Uremic Toxic Blood-Brain Barrier Disruption Mediated by AhR Activation Leads to Cognitive Impairment during Experimental Renal Dysfunction[J]. Journal of the American Society of Nephrology, 2020, 31(7): 1509.

[7] Chen H J, Wang Y F, Qi R, et al. Altered Amygdala Resting-State Functional Connectivity in Maintenance Hemodialysis End-Stage Renal Disease Patients with Depressive Mood[J]. Molecular Neurobiology, 2017, 54(3): 2223–2233.

[8] Li A, Mu J, Huang M, et al. Altered amygdala-related structural covariance and resting-state functional connectivity in end-stage renal disease patients[J]. Metabolic Brain Disease, 2018, 33(5): 1471–1481.

[9] Del Prete D, Gambaro G, Lupo A, et al. Precocious activation of genes of the renin-angiotensin system and the fibrogenic cascade in IgA glomerulonephritis[J]. Kidney International, 2003, 64(1): 149–159.

[10] Gong S, Deng F. Renin-angiotensin system: The underlying mechanisms and promising therapeutical target for depression and anxiety[J]. Frontiers in Immunology, 2023, 13: 1053136.

[11] Kloet A D de, Wang L, Pitra S, et al. A Unique “Angiotensin-Sensitive” Neuronal Population Coordinates Neuroendocrine, Cardiovascular, and Behavioral Responses to Stress[J]. Journal of Neuroscience, 2017, 37(13): 3478–3490.

[12] Faghih M, Drewes A M, Singh V K. Psychiatric Disease Susceptibility and Pain in Chronic Pancreatitis: Association or Causation?[J]. The American Journal of Gastroenterology, 2021, 116(10): 2026–2028.

[13] Dantzer R, O’Connor J C, Freund G G, et al. From inflammation to sickness and depression: when the immune system subjugates the brain[J]. Nature Reviews. Neuroscience, 2008, 9(1): 46–56.

[14] Cao W, Yang Z, Liu X, et al. A kidney-brain neural circuit drives progressive kidney damage and heart failure[J]. Signal Transduction and Targeted Therapy, 2023, 8(1): 1–11.

[15] Tan R Z, Zhong X, Li J C, et al. An optimized 5/6 nephrectomy mouse model based on unilateral kidney ligation and its application in renal fibrosis research[J]. Renal Failure, 2019, 41(1): 555–566.

[16] Motohashi H, Tahara Y, Whittaker D S, et al. The circadian clock is disrupted in mice with adenine-induced tubulointerstitial nephropathy[J]. Kidney International, 2020, 97(4): 728–740.

[17] Peng H, Wang Q, Lou T, et al. Myokine mediated muscle-kidney crosstalk suppresses metabolic reprogramming and fibrosis in damaged kidneys[J]. Nature Communications, 2017, 8(1): 1493.

[18] Li X Q, Jin B, Liu S X, et al. Neddylation of RhoA impairs its protein degradation and promotes renal interstitial fibrosis progression in diabetic nephropathy[J]. Acta Pharmacologica Sinica, 2025, 46(6): 1692–1705.

[19] Shao J, Chen Y, Gao D, et al. Ventromedial hypothalamus relays chronic stress inputs and exerts bidirectional regulation on anxiety state and related sympathetic activity[J]. Frontiers in Cellular Neuroscience, 2023, 17: 1281919.

[20] Wu X, Xu W, Deng L, et al. Spatial multi-omics at subcellular resolution via high-throughput in situ pairwise sequencing[J]. Nature Biomedical Engineering, 2024, 8(7): 872–889.

[21] Villapol S, Yaszemski A K, Logan T T, et al. Candesartan, an Angiotensin II AT1-Receptor Blocker and PPAR-γ Agonist, Reduces Lesion Volume and Improves Motor and Memory Function After Traumatic Brain Injury in Mice[J]. Neuropsychopharmacology, 2012, 37(13): 2817–2829.

[22] Kidokoro K, Cherney D Z I, Bozovic A, et al. Evaluation of Glomerular Hemodynamic Function by Empagliflozin in Diabetic Mice Using In Vivo Imaging[J]. Circulation, 2019, 140(4): 303–315.

[23] Hochgerner H, Singh S, Tibi M, et al. Neuronal types in the mouse amygdala and their transcriptional response to fear conditioning[J]. Nature Neuroscience, 2023, 26(12): 2237–2249.

[24] Yu Y H, Kim S W, Park D K, et al. Altered Emotional Phenotypes in Chronic Kidney Disease Following 5/6 Nephrectomy[J]. Brain Sciences, 2021, 11(7): 882.

[25] Renczés E, Marônek M, Gaál Kovalčíková A, et al. Behavioral Changes During Development of Chronic Kidney Disease in Rats[J]. Frontiers in Medicine, 2020, 6: 311.

[26] Mazumder M K, Giri A, Kumar S, et al. A highly reproducible mice model of chronic kidney disease: Evidences of behavioural abnormalities and blood-brain barrier disruption[J]. Life Sciences, 2016, 161: 27–36.

[27] Manigault A W, Kuhlman K R, Irwin M R, et al. Psychosocial resilience to inflammation-associated depression: a prospective study of breast-cancer survivors[J]. Psychological Science, 2022, 33(8): 1328–1339.

[28] Manigault A W, Kuhlman K R, Irwin M R, et al. Vulnerability to inflammation-related depressive symptoms: moderation by stress in women with breast cancer[J]. Brain, Behavior and Immunity, 2021, 94: 71–78.

[29] Babaev O, Piletti Chatain C, Krueger-Burg D. Inhibition in the amygdala anxiety circuitry[J]. Experimental & Molecular Medicine, 2018, 50(4): 1–16.

[30] Fadok J P, Krabbe S, Markovic M, et al. A competitive inhibitory circuit for selection of active and passive fear responses[J]. Nature, 2017, 542(7639): 96–100.

[31] Saavedra J M, Sánchez-Lemus E, Benicky J. Blockade of brain angiotensin II AT1 receptors ameliorates stress, anxiety, brain inflammation and ischemia: therapeutic implications[J]. Psychoneuroendocrinology, 2011, 36(1): 1–18.

[32] Benicky J, Sánchez-Lemus E, Honda M, et al. Angiotensin II AT1 Receptor Blockade Ameliorates Brain Inflammation[J]. Neuropsychopharmacology, 2011, 36(4): 857–870.

[33] Costa-Ferreira W, Morais-Silva G, Gomes-de-Souza L, et al. The AT1 receptor antagonist losartan does not affect depressive-like state and memory impairment evoked by chronic stressors in rats[J]. Frontiers in Pharmacology, 2019, 10: 705.

[34] Badoer E. Role of the hypothalamic PVN in the regulation of renal sympathetic nerve activity and blood flow during hyperthermia and in heart failure[J]. American Journal of Physiology Renal Physiology, 2010, 298(4): F839–F846.

[35] Liu Y, Rao B, Li S, et al. Distinct hypothalamic paraventricular nucleus inputs to the cingulate cortex and paraventricular thalamic nucleus modulate anxiety and arousal[J]. Frontiers in Pharmacology, 2022, 13: 814623.

