## Supplementary figures and images for "Chronic kidney disease promotes anxiety susceptibility through an angiotensin II–central amygdala axis"

### figure S1

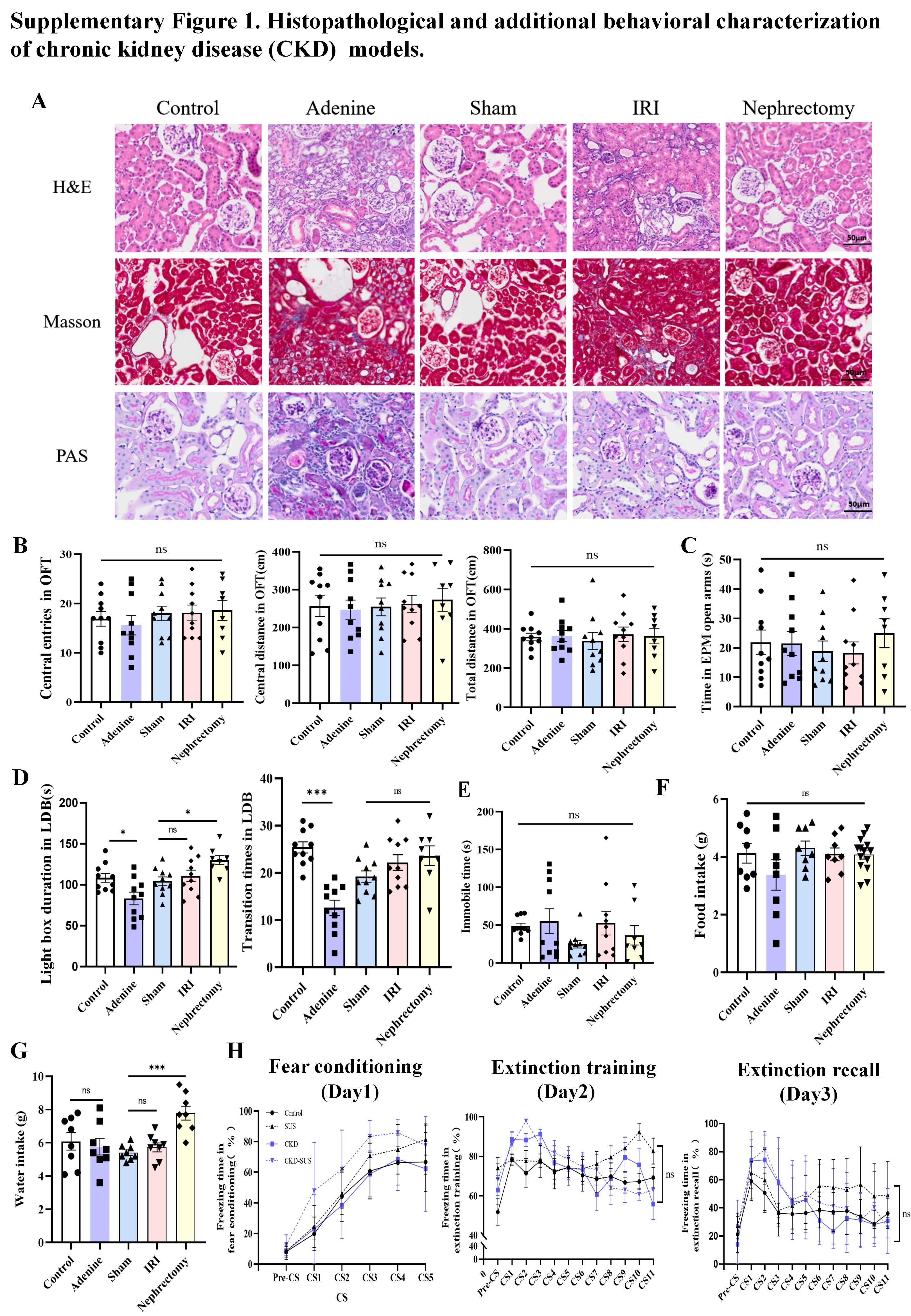

### figure S2

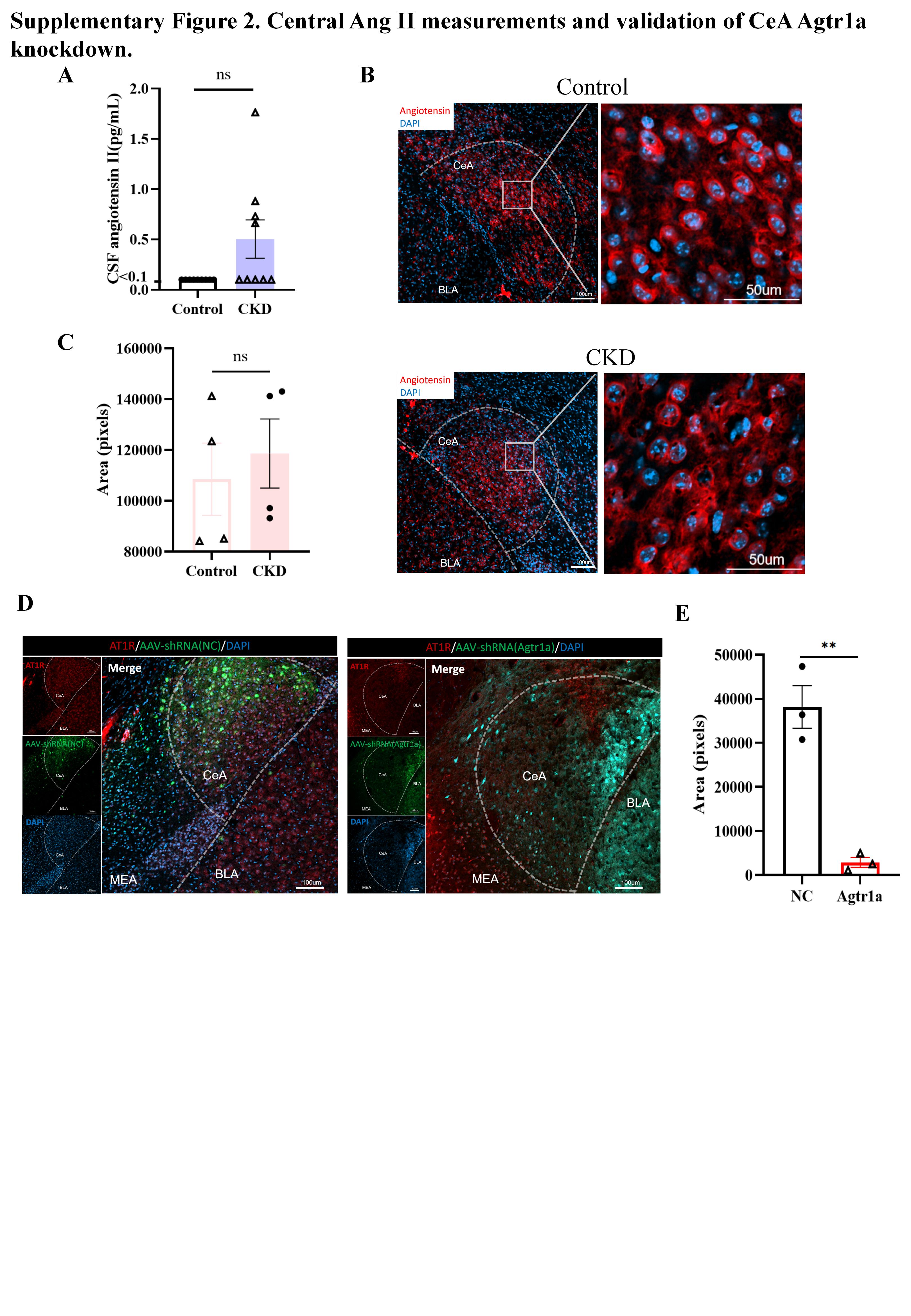

### figure S3

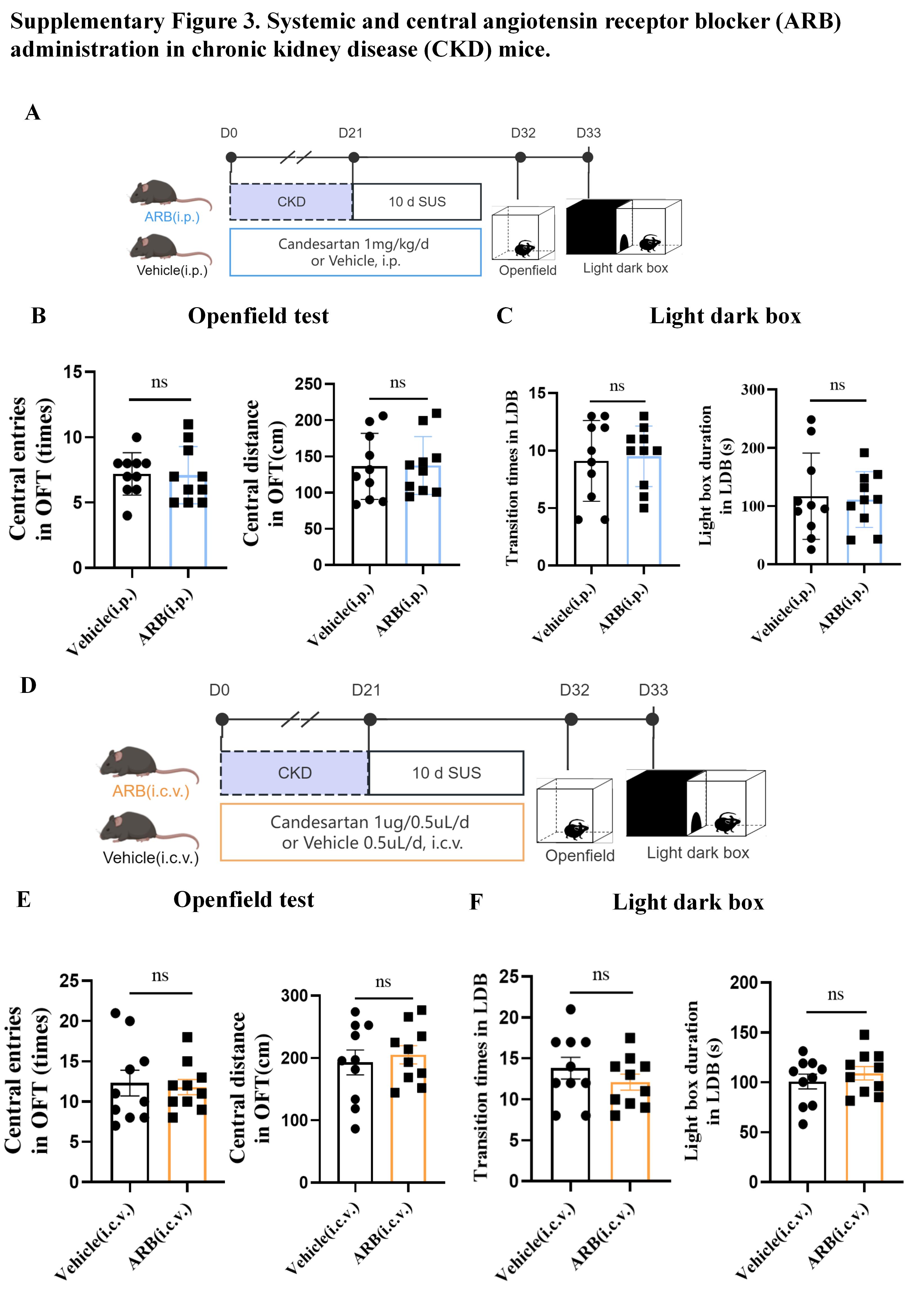
